# Viscoelastic relaxation of collagen networks provides a self-generated directional cue during collective migration

**DOI:** 10.1101/2020.07.11.198739

**Authors:** Andrew G. Clark, Ananyo Maitra, Cécile Jacques, Anthony Simon, Carlos Pérez-González, Xavier Trepat, Raphaël Voituriez, Danijela Matic Vignjevic

## Abstract

There is growing evidence that the physical properties of the cellular environment can impact cell migration. However, it is not currently understood how active physical remodeling of the network by cells affects their migration dynamics. Here, we study collective migration of small clusters of cells on deformable collagen-1 networks. Combining theory and experiments, we find that cell clusters, despite displaying no apparent internal polarity, migrate persistently and generate asymmetric collagen gradients during migration. We find that persistent migration can arise from viscoelastic relaxation of collagen networks, and reducing the viscoelastic relaxation time by chemical crosslinking leads to a reduction in migration persistence. Single cells produce only short range network deformations that relax on shorter timescales, which leads to lower migration persistence. This physical model provides a mechanism for self-generated directional migration on viscoelastic substrates in the absence of internal biochemical cues.

## Introduction

Collective cell migration is an essential process during development and tissue homeostasis and has also been proposed to play a role in early stages of cancer metastasis^1, 2^. During invasive stages of epithelial cancers, which comprise 80-90% of human cancers^3^, cells or groups of cells migrate through the surrounding stroma, which is comprised primarily of collagen-1 extracellular matrix (ECM) networks. It has been proposed that cells migrating in the stroma follow chemotactic cues to reach vessels, where they can then intravasate and be transported to secondary organs^4^.

In addition to chemical cues, collagen fiber thickness and alignment can also act as a directional cue during cell migration^5–7^. Large cell aggregates embedded in collagen networks *in vitro* have been shown to physically pull on collagen fibers, resulting in radially aligned collagen bundles that facilitate invasion of single cells into the surrounding matrix^8–10^. Such reorganization of stromal collagen networks has been associated with cancer invasiveness, and radial arrays of thick collagen bundles have been proposed to act as “highways” for tumor dissemination^11, 12^. However, collagen reorganization and cell migration in these systems occur on drastically different length- and timescales, and by the time cells begin to migrate, the collagen network has already been reorganized. These studies clearly show that existing collagen topologies can direct cell migration, but they do not account for the active reorganization of collagen networks by cells during migration.

In this study, we combined long-term imaging, traction force microscopy and theoretical modeling to ask how the viscoelastic properties of collagen networks affect network reorganization during collective cell migration and how these local changes in network topology can feed back on migration dynamics. Our results suggest that viscoelastic relaxation in collagen gels can give rise to local asymmetries in collagen organization that drive spontaneous persistent migration, even in the absence of biochemical polarity cues.

## Results

### Cell clusters migrate more persistently on collagen networks compared to soft elastic gels

We first sought to understand how collective cell migration dynamics differed between migration on deformable collagen networks vs. soft elastic substrates. To this end, we generated small (2-50 cell) clusters of A431 cells, which have previously been shown to migrate collectively as small clusters on collagen/Matrigel composite networks^13^. We cultured A431 clusters on soft (0.5kPa) poly-A-acrylamide (PAA) gels coated with either a thin (∼30*µ*m) layer of polymerized Collagen-1 (2mg/ml) or non-polymerized monomeric Collagen-1 and tracked cell migration over ∼16h (Figure 1a, Supplementary Movies 1,2). Clusters appeared to migrate along straighter tracks and explore a larger area when migrating on collagen networks compared to monomeric collagen.

**Figure 1.**
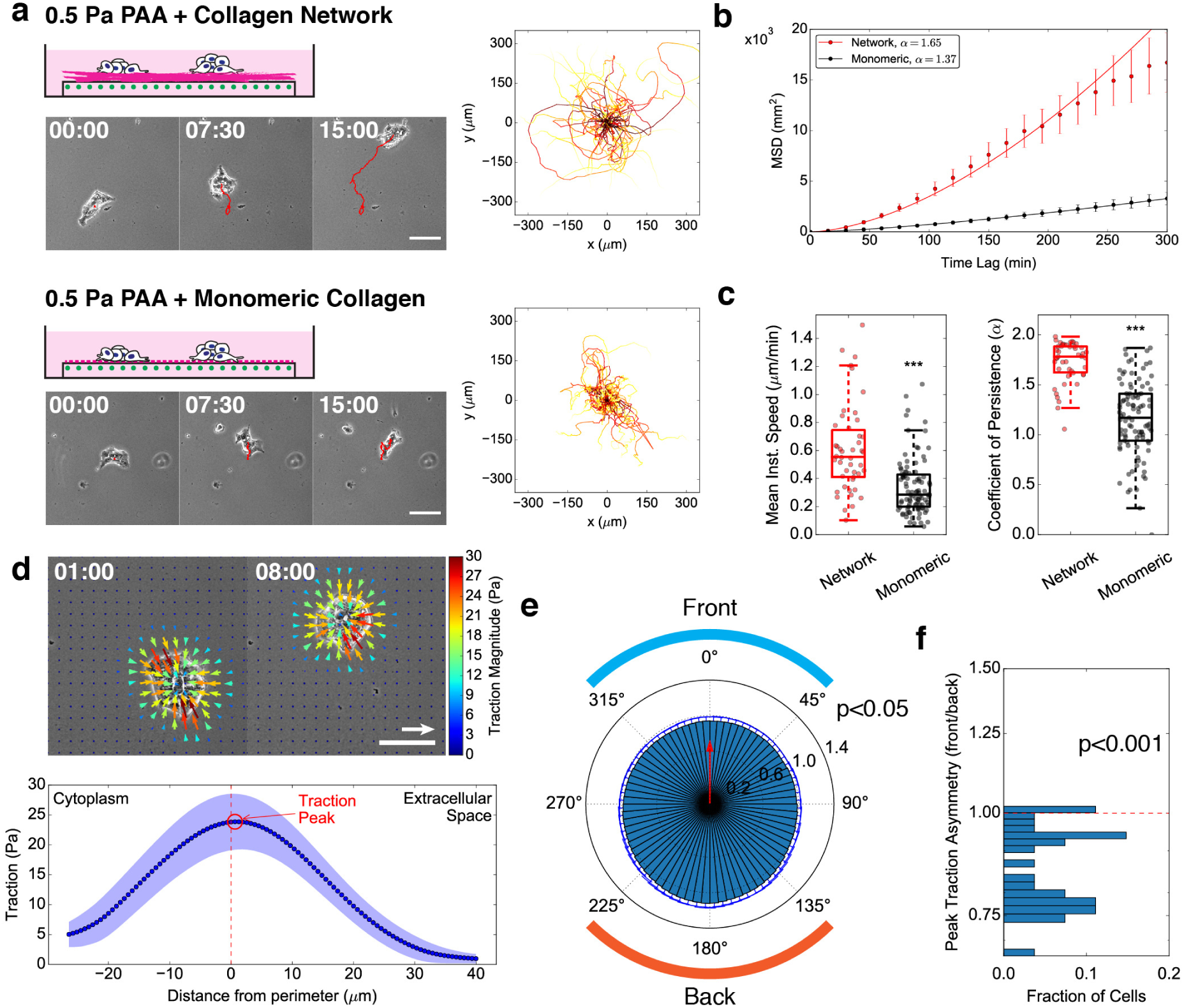
Cell clusters exhibit persistent migration on collagen networks with asymmetric traction force distributions. **a.** *Left:* Schematics and montages of A431 cell clusters migrating on 0.5 kPa PAA gels coated with a thin collagen-1 network (*top panels*) or 100*µ*g/ml monomeric collagen-1 (*bottom panels*, see also Supplementary Movies 1,2). Scale Bar: 100*µ*m. HH:MM. *Right:* Overlaid cluster migration trajectories, adjusted so that all trajectories start at the origin (0,0). **b.** Mean squared displacement (MSD) curves for cell clusters migrating on PAA gels coated with collagen network (red) or monomeric collagen (black). **c.** Boxplots of mean instantaneous speed (*left*) and coefficient of persistence (*α, right*). Coefficient of persistence was determined by a linear fit to the logarithm of the MSD curve for each trajectory. Each dot represents one cluster trajectory. For *b* and *c*, n = 46, 97 from N = 5, 3 independent experiments. ***p<0.001 for Welch’s t-test. **d.** *Top:* Example micrograph overlaid with traction force vectors (arrows) from traction force microscopy experiment of cluster migration on collagen network on a PAA gel (see also Supplementary Movie 4). Scale bar: 100*µ*m. Scale vector: 50Pa. HH:MM. *Bottom:* Average linescan (mean±SD) of traction forces around the cell cluster in the upper panel. Red dotted line: the cluster periphery. Red circle: peak traction force magnitude. **e.** Polar plot of peak traction force magnitude with respect to migration direction (red arrow). Values are mean±SD, averaged over all timepoints for n = 27 clusters from N = 3 independent experiments. **f.** Histogram of peak traction force asymmetry (mean of the front quadrant divided by the mean of the rear quadrant, from the same data averaged in *e*). The y-axis is shown in log-scale. P-value reflects t-test with *µ*_0_ = 1.

To determine whether clusters migrated with higher persistence (i.e. along straighter paths), we calculated the average mean squared displacement (MSD; Figure 1b). Fitting the MSD curves with a power-law function, we extracted a higher exponent (the “coefficient of persistence”, *α*) for clusters migrating on collagen networks, suggesting that clusters migrate more persistently on collagen networks. We then extracted the mean instantaneous speed and a coefficient of persistence for individual trajectories (Figure 1c). We found that clusters on collagen networks migrated faster and more persistently than clusters migrating on collagen-coated PAA gels. We found no significant difference in migration dynamics on PAA gels coated with different concentrations of monomeric collagen-1, suggesting that ligand density cannot account for the difference in migration on collagen networks vs. monomeric collagen (Figure S1a,b). These data suggest that cell clusters migrate faster and more persistently on collagen networks compared to collagen-coated elastic gels.

### Cell clusters migrating on collagen networks do not display canonical front-back polarity

One potential explanation for persistent collective migration on collagen networks could be front-back polarity mechanisms at the cluster scale. To investigate this, we first immunostained cell clusters on collagen networks and stained for Rac1, a prominent front-back marker that is typically associated with the protrusive leading edge during single cell migration^14^. We found that Rac1 is localized at the cell cortex specifically at the periphery of the clusters; Rac1 appears to be down-regulated at cell-cell junctions at the cluster interior (Figure S2a). This is consistent with a previous report showing that Rho family GTPases and myosin-2 are down-regulated at interior junctions by the DDR1-Par3/6-RhoE pathway^13^. This suggests that cell clusters behave as large “super cells”.

Front-back polarization is a common mechanism by which cells migrate persistently. Myosin-2 is typically found at the cell rear in front-back polarized cells and is involved in symmetry breaking and polarized cell migration^14–16^. In order to determine whether myosin-2 was polarized during persistent cluster migration on collagen networks, we performed live imaging of a stable A431 cell line expressing a myosin-2 light chain fused to green fluorescent protein (A431 MLC-GFP). We then segmented the cortical and interior regions of each cluster over time and compared the direction of migration with the relative cortical intensity of MLC-GFP around the cluster perimeter (Figure S2b, Supplementary Movie 3). We found no asymmetry in the cortical MLC signal with respect to the migration direction. We also did not observe any preferential orientation in centrosome position with respect to the nuclei (Figure S2c) or shape asymmetries associated with migration (Figure S3). Together, these data suggest that cell clusters do not display typical front-back polarity mechanisms during collective migration on collagen networks.

### Cell clusters migrating on collagen networks generate asymmetric traction force patterns

We next sought to determine whether cell clusters exhibit asymmetric traction forces during migration, which could provide a physical mechanism for persistent migration on collagen networks. To this end, we performed traction force microscopy (TFM) on clusters migrating on thin collagen networks polymerized on soft PAA gels (Figure 1d, *upper panels*; Supplementary Movie 4). Because tractions are measured at the surface of the PAA gels, they provide information on how cell-generated forces are transmitted through the thin collagen layer. We found that cell clusters generated radial inward-facing tractions around the perimeter of the cluster. To better assess the magnitude of traction forces with respect to the periphery of the cluster, where the contractile cortical components localized, we segmented the cells and generated traction force linescans from the center of the cluster to 40*µ*m outside of the cluster (Figure 1d, *lower panel*). As consequence of the thin collagen layer, tractions peaked slightly outside the cluster contour, rather than right at the contour as usually observed when clusters are in direct contact with the PAA gel^17, 18^.

To determine whether the distribution of traction forces was asymmetric during cluster migration, we analyzed traction force linescans along the periphery of the clutster (Figure S4a). We found that the peak in traction magnitude was highest at the cluster rear. To further examine the difference in the traction force pattern at the cluster front and rear, we projected the tractions along the axis of migration (Figure S4b). We found that projected tractions were negative at the front, where tractions were rearward-facing, and positive at the rear, where traction forces were forward-facing. Taking linescans of the projected tractions from the front to the rear of the cluster (Figure S4c,d), we found that the traction peak was slightly higher in the rear, while the tail of traction decay at the front was slightly wider, indicating a slight asymmetry in the distribution of traction forces at the front vs. rear of the cluster. As expected from force balance, the vectorial sum of the tractions was zero.

To quantify the asymmetry in traction force distributions, we measured the peak traction magnitude with respect to the migration direction, averaged over time and over different clusters (Figure 1e). We found that the traction peak at the rear of the cluster was ∼10% higher compared to the leading edge of the cluster (Figure 1f). These data suggest that cell clusters induce slightly asymmetric traction force profiles on collagen networks during migration.

### Local collagen topology is asymmetric during collective migration

We hypothesized that the asymmetric traction pattern would correlate with an asymmetric deformation of the collagen network on which these clusters migrate. To investigate this, we imaged migration of cell clusters on fluorescently-labeled collagen networks (Figure 2a). We observed that cell clusters reorganized collagen networks during migration, generating radial arrays of collagen fibers around the cluster, as would be expected from the inward facing tractions around the periphery (Figure 1d). To assess the degree of collagen alignment around clusters, we extracted local filament orientations of fibers on the networks surface (Figure S5a,b). The average nematic order outside of the cluster was ∼0.6, indicating that the filaments were indeed aligned in a radial pattern around the clusters, though the degree of filament orientation was symmetric around the outside of the cluster (Figure S5c).

**Figure 2.**
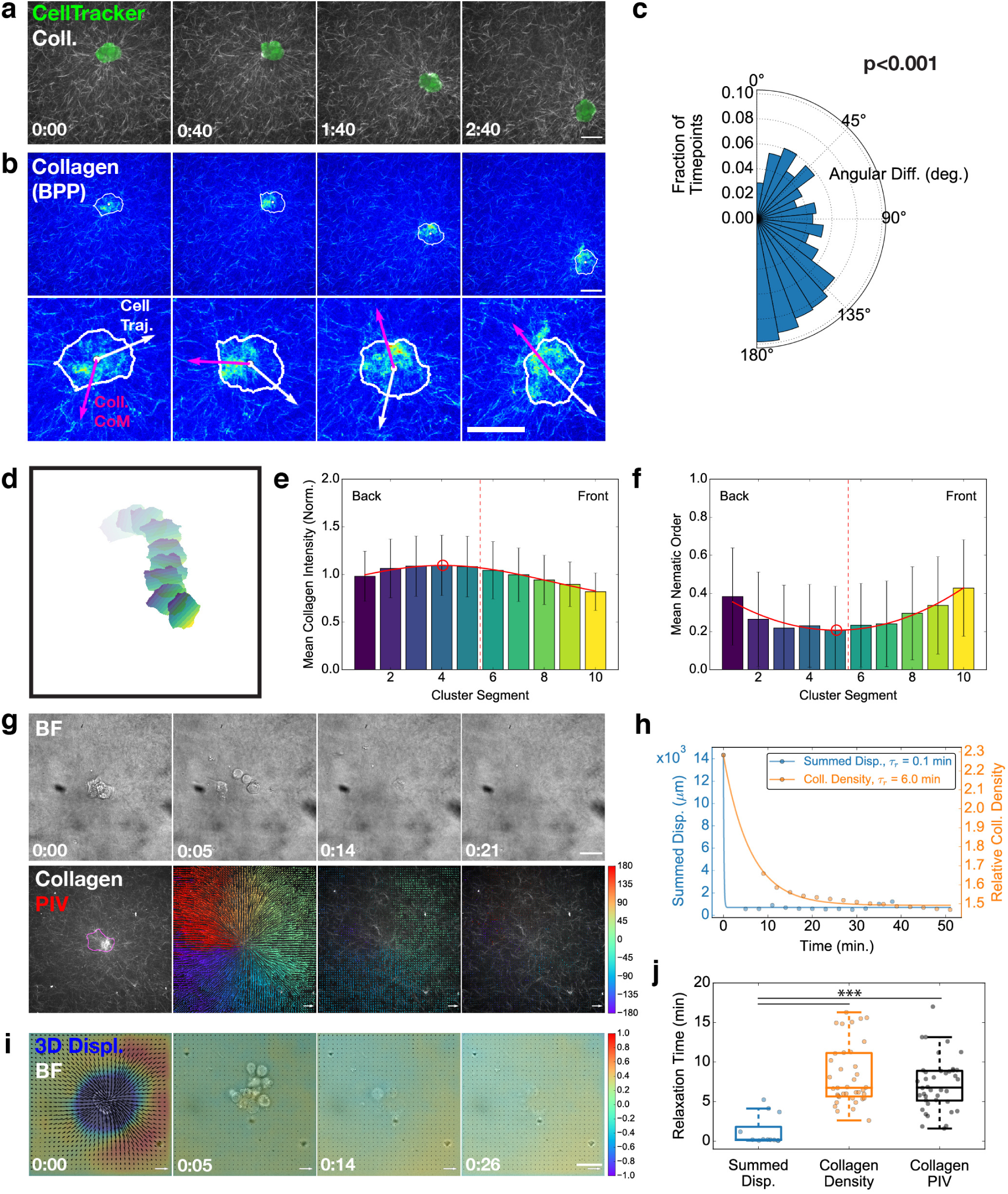
Local collagen topology is asymmetric during collective migration and relaxes viscoelastically. **a.** Montage of a cluster migrating on a fluorescent collagen network. Green: brightest point projection of cluster labeled with CellTracker Green. Grayscale: single *z*-slice of fluorescent collagen surface. Scale Bar: 50*µ*m. HH:MM. **b.** Pseudocolor montage of fluorescent collagen (brightest point projection; see also Supplementary Movie 5). White line: segmented cluster contour. White dot: center of mass of the cluster area. *Lower panel*: zoom of cluster from upper panel. White arrow: vector of cell migration trajectory. Magenta arrow: vector from center of mass of cluster to weighted center of mass of collagen. Scale Bar: 25*µ*m. HH:MM. **c.** Polar histogram of angular difference between trajectory and collagen center of mass vectors for all timepoints. P-value reflects Rayleigh test of circularity. **d.** Overlaid segmentations from the time series shown in *a*, split into 10 regions from the front (yellow) to the rear (violet) for each time point. **e.** Bar plot of the collagen intensity over the cluster regions (mean±SD). **f.** Bar plot of the collagen fiber nematic order over the cluster regions (mean±SD). For panels *c,e* and *f*, data was averaged over all timepoints for n = 34 clusters from N = 5 independent experiments. **g.** Montage of brightfield and fluroescent collagen signal (brightest point projection) with vectors from PIV analysis overlaid (see also Supplementary Movie 6). Magenta: segmented cell contour. Immediately following t=0:00, cells were removed using Trypsin/NH4OH and imaging was resumed. HH:MM. **h.** Time course of summed 3D displacements (blue) and collagen density within the cell contour (orange) for the experiments in *g* and *i*. Dots: data from individual time points. Line: fit of the data with an exponential decay function to extract the average relaxation time, *τ*_*r*_. **i.** Montage of 3D displacements of a cell cluster on a PAA gel coated with a collagen network, with cell removal after 0:00 (see also Supplementary Movie 4). Black arrows: *xy* displacements. Color scale: *z* displacements. Negative values indicate downward displacement (toward the collagen/PAA). HH:MM. **j.** Boxplot of relaxation times extracted from 3D substrate displacements, collagen density measurements and collagen recoil by PIV. Each dot represents a cluster that was rapidly removed. One-way ANOVA p<0.001. ***p<0.001 for post-hoc Tukey HSD test. Data represents n = 12, 39 clusters from N = 3, 6 independent experiments (3D displacement, Collagen relaxation, respectively).

We next investigated the local collagen topology in the region directly underlying the clusters, which could affect migration dynamics via direct contact with the cell clusters. We segmented cells and analyzed the fluorescent collagen signal in the segmented region (Figure 2b, *upper panels*; Supplementary Movie 5). It appeared from these images that collagen density was asymmetrically patterned; a region of high collagen density was typically observed between the center of mass and the trailing edge of the cell cluster. To quantify this, we defined a vector pointing from the cluster center of mass to the weighted center of mass of the collagen signal inside of the cluster boundaries (Figure 2b, *lower panels, magenta arrows*). We compared the angle of this vector to the angle of the cell displacement (Figure 2b, *lower panels, white arrows*). We found that the angular difference between these vectors was asymmetrically distributed, with a higher probability toward angular differences of 180°C, indicating that cells tend to move away from regions of high collagen density (Figure 2c). Thus, simply by observing the cell position and underlying collagen density, we can predict the most probable direction of cluster migration.

To further investigate how collagen density varied along the length of the cluster, we split the segmentation into ten regions of equal length from the front to the rear of the cluster (Figure 2d). Quantifying the average collagen density in each region over all timepoints and clusters, we found that collagen density reached a maximum peak behind the cluster center (Figure 2e). We then measured the nematic order of filaments in each region with respect to the direction of migration. We found that filament orientation reached a minimum behind the cluster center (Figure 2f). These data suggest that clusters generate inverse patterns of collagen density and nematic order during cluster migration, with a peak that is offset toward the trailing edge of the cluster.

### Collagen networks behave viscoelastically in response to tractions generated by cell clusters

We hypothesized that the offset between the peak of collagen density/orientation and the center of the cluster could be due to a slow, viscous-like relaxation of the collagen network during migration. Collagen networks have indeed been shown to relax in viscoelastic manner when applied stress is released^19^. To investigate viscoelastic behavior in collagen networks in response to tractions forces generated by cell clusters, we rapidly removed clusters by acute treatment with Trypsin and Ammonium Hydroxide (NH_4_OH; Figure 2g, Supplementary Movie 6). To determine how the collagen networks responded to rapid cell removal, we tracked the collagen density in the segmented cluster region over time and fit the decay with an exponential function to determine a relaxation timescale, *τ*_*r*_ (Figure 2h). The relative collagen density relaxed to a value greater than one (1.78±0.66; mean±SD), suggesting that these collagen networks are, to some degree, plastic, consistent with a previous report showing that cell-scale forces can permanently remodel collagen networks^20^. To further characterize the collagen relaxation, we performed particle image velocimetry (PIV) analysis on the collagen images to track frame-to-frame movements in the collagen fibers during relaxation (Figure 2g, *lower panels*).

To ensure that Trypsin/NH_4_OH treatment indeed resulted in an immediate removal of stress induced by cell clusters on the collagen networks, we performed 3D displacement microscopy. In 3D displacement microscopy, clusters are plated on PAA gels with fluorescent beads coated with collagen networks. The deformation of the substrate, which is caused by physical stresses generated by the cluster, is determined by measuring the 3D bead positions relative to a reference image. Prior to the Trypsin/NH_4_OH treatment, cells exerted radial, inward facing stresses in *x* and *y*, consistent with our 2D TFM results, as well as downward facing stresses in *z*. Such downward facing stresses have been previously reported during *Dictyostelium* migration^21^. Immediately upon cell removal, the substrate displacements disappeared (Figure 2i, Supplementary Movie 7). We measured the summed 3D displacements over time and extracted the relaxation time (Figure 2h). We found that the relaxation time of the displacements was significantly smaller than the relaxation time of collagen density or collagen movements measured by PIV (Figure 2j). Together, these data suggest that collagen networks behave in a viscoelastic manner in response to stresses generated by cell clusters during migration.

### Theoretical model of persistent migration on a viscoelastic substrate

To investigate whether the asymmetric collagen distributions generated during migration could facilitate persistent migration, we developed a simple theoretical model to identify the minimal ingredients required for persistent polarized migration of a cell cluster without any intrinsic polarity or shape asymmetry (see Supplementary Note). The main ingredient of the model is the active coupling of an isotropic cluster to an apolar viscoelastic medium, which we show is sufficient to induce symmetry-breaking and thereby persistent motion of the cluster.

The model describes the cell cluster as an isotropic active particle, which can be assumed to be point-like, and whose position along a one dimensional axis is denoted by *x*_*c*_; its velocity is denoted by *v*_*c*_. This active particle deforms the viscoelastic substrate, causing a structural perturbation *S*(*x, t*) (Figure 3a), whose equation of motion can be schematically written as

**Figure 3.**
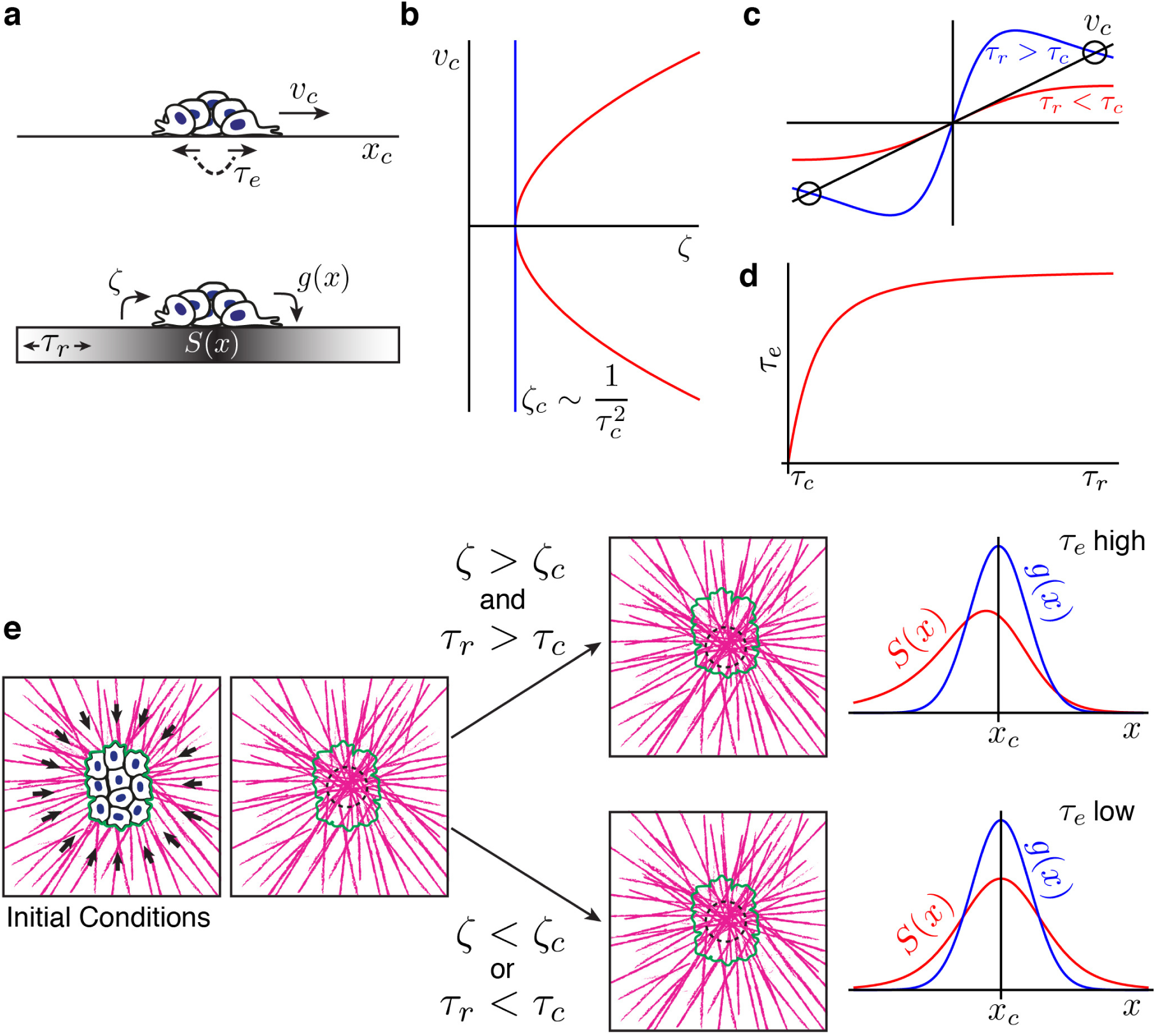
Theoretical model of persistent migration on a viscoelastic substrate. **a.** Schematic description of the model. A cluster of cells moves along the axis *x*_*c*_ with velocity *v*_*c*_. The probability of switching migration direction is given by the timescale *τ*_*e*_. The cell cluster interacts with the substrate according to a symmetric forcing function *g*(*x*), resulting in a deformation of the substrate given by *S*(*x*). The deformation in the substrate leads to activity in the cell cluster via the coupling *ζ*. As the substrate is viscoelastic, the relaxation is characterized by the timescale *τ*_*r*_. **b.** The cell migration velocity *v*_*c*_ undergoes a supercritical pitchfork bifurcation at *ζ* = *ζ*_*c*_. Here, *v*_*c*_ is plotted as a function of *ζ* for *g*(*ξ*^*′*^) = *e*^−*ξ*′2/2^. **c.** Solutions for migration velocity *v*_*c*_ for substrate relaxation times above or below the critical relaxation time. Solutions are shown by the intersection with the black line. While for *τ*_*r*_ < *τ*_*c*_, only a stable (*v*_*c*_ = 0) solution exists, a positive solution for *v*_0_ exists for *τ*_*r*_ > *τ*_*c*_. **d.** Cluster persistence time scale *τ*_*e*_ as a function of the substrate relaxation time scale *τ*_*r*_ relative to the critical relaxation time *τ*_*c*_. **e.** Interpretation of the model as it relates to experiments. Initially symmetric inward-directed tractions generate radially oriented collagen fibers, with a high collagen density in the center of the cluster. After symmetry breaking, for networks with sufficient cell-substrate coupling (*ζ* > *ζ*_*c*_) and relaxation time (*τ*_*r*_ > *τ*_*c*_), clusters are predicted to migrate persistently (i.e. *τ*_*e*_ is high) due to the offset between the driving function *g*(*x*) and the substrate deformation *S*(*x*). If the coupling or relaxation times are below the critical values, *g*(*x*) and *S*(*x*) are both symmetric around *x*_*c*_ and low persistence migration (low *τ*_*e*_) is predicted.

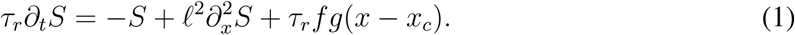

Here, *τ*_*r*_ is the viscoelastic relaxation time of the collagen network, *l* is a deformation lengthscale over which a point perturbation spreads out, *g*(*x* − *x*_*c*_) is a normalized function which encodes the functional form of the source of the perturbation of the collagen network due to the active particle and is expected to peak at *x*_*c*_ and vanish for *x* − *x*_*c*_ ≫ 0, and *f* is the strength of the perturbation (proportional to the total stress exerted by the cluster on the network). This is distinct from previous continuum models of cell-substrate interactions^22–24^.

The nature of the perturbation is not specific and could represent, for example, a local change in substrate density and/or the scalar nematic order parameter, both of which we observe experimentally (Figure 2e,f). Importantly, the source of the perturbation *g* is assumed to be spatially symmetric, meaning that the cluster alone does not carry any internal polarity. In the cases of perturbations of density and nematic order, this phenomenological equation, derived explicitly in the Supplementary Note, assumes a minimal rheological description of the collagen gel as a linear elastic solid (of modulus *E*) with viscous relaxation (of viscosity *η* ∝ *Eτ*_*r*_). The isotropic particle actively responds to the substrate perturbation via a phenomenological coupling *ζ*, which parameterizes the active response of the cluster to the perturbation: 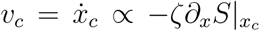. Although the precise biological source(s) of the active coupling parameter *ζ* is not known, this coupling could, for example, result in the asymmetric traction forces that we observe during collective migration (Figure 1e,f).

This simple model predicts that at a critical value of coupling *ζ*_*c*_, the particle’s velocity will display a supercritical pitchfork bifurcation (Figure 3b). For *ζ* < *ζ*_*c*_, the only solution is *v*_*c*_ = 0, and the cluster cannot migrate persistently. However, for *ζ* > *ζ*_*c*_, one finds *v*_*c*_ ∼ ±(*ζ* −*ζ*_*c*_)^1/2^, implying that the cluster migrates persistently in a random direction. Importantly, the critical value of activity scales with the relaxation time of the substrate as 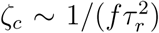. This implies that increasing the relaxation time of the collagen network beyond a critical value *τ*_*c*_ at a fixed value of the coupling *ζ* leads to a persistent motion of the cluster. It is likely that the intensity *f* of the perturbation would depend on cluster size *L*, as smaller clusters or single cells are known to produce lower traction force^17^. In addition, *ζ* is also likely to depend on cell size, as clusters that are small compared to the deformation length scale *l* would be too small to sense a collagen gradient. Persistent motion is therefore predicted to occur only for large enough clusters on substrates with long enough relaxation times. In the presence of noise, the model predicts a Brownian-like random motion for *ζ* < *ζ*_*c*_ or *τ*_*r*_ < *τ*_*c*_. For *ζ* > *ζ*_*c*_ at fixed *τ*_*r*_ or *τ*_*r*_ > *τ*_*c*_ at fixed *ζ*, the model predicts persistent motion, whose persistence time increases with *τ*_*r*_ (Figure 3d).

In the model, persistent migration arises from the interaction of the moving cluster with the viscoelastic substrate (Figure 3e). The activity of the cluster induces a perturbation in the substrate, analogous to traction forces causing local changes in filament density/orientation. For a stationary cluster, both the force from the cluster and the perturbation of the substrate have symmetric profiles around the cluster center, which does not cause any motion. Let us now consider a cluster moving at speed *v*_*c*_. Because of the viscoelastic relaxation of the substrate, the cluster position is *ahead* of the peak of the substrate deformation profile by *d* ∼ *v*_*c*_*τ*_*r*_. This implies that the cluster experiences an active force due to the substrate asymmetry ∝ −*∂*_*x*_*S*, which increases with *d*, making it slide downhill along the perturbation profile (assuming *ζ* > 0 without loss of generality). For small *τ*_*r*_, the active force is not sufficient to sustain the speed *v*_*c*_ and the cluster velocity relaxes to 0; however, for large enough *τ*_*r*_ the active force is sufficient to sustain the steady motion at *v*_*c*_ and the particle can surf the deformation profile (which it itself induces) at a constant speed, leading to persistent migration. The model qualitatively recapitulates our initial observations and predicts an offset between the cluster center and deformation peak on the order of the product of the cell velocity and relaxation time (*d* ∼ *v*_*c*_*τ*_*r*_). With *v*_*c*_ ≈ 0.6*µ*m/min (Figure 1c) and *τ*_*r*_ ≈ 8min (Figure 2j), we therefore expect *d* ∼ 5*µ*m. From the average gradient data, the average rearward offset of the collagen density peak is 9.0*µ*m, and the average rearward offset of the nematic order trough is 2.9*µ*m, in line with the predictions of the theory.

Our model presents a theoretical picture of how a statistically isotropic cell cluster may nevertheless move persistently due to the anisotropy it induces in the substrate. Motion occurs via a spontaneous symmetry-breaking mechanism that is conceptually similar to a model proposed for autophoretic colloids^25^. The theory makes two additional important predictions: (1) migration persistence should decrease for substrates with lower relaxation times and (2) migration persistence should be lower for small clusters. In the Supplementary Note we further provide a general theoretical framework that describes the active dynamics of an active system localised in space (the cluster), which interacts with a generic viscoelastic nematic substrate. The analysis of this more general description shows that the above mechanism leading to persistent motion can be generalized and could be at work in other active systems, living or artificial.

### Collagen crosslinking decreases viscous relaxation time and reduces persistent migration

We first sought to test the prediction that reducing substrate relaxation time leads to reduced persistence. To this end, we compared control collagen networks with collagen networks treated with the small sugar threose, which has previously been shown to crosslink collagen networks^10^. We analyzed collagen density and relaxation following rapid cell removal (Figure 4a, Supplementary Movies 8-10). Crosslinking with threose led to reduced initial collagen density prior to cluster removal, suggesting an increase in network stiffness as previously observed^10^. Crosslinking with threose also led to a reduction in relaxation time following cluster removal (Figure 4b). These data suggest that threose crosslinking leads to stiffer networks that relax faster compared to noncrosslinked networks.

**Figure 4.**
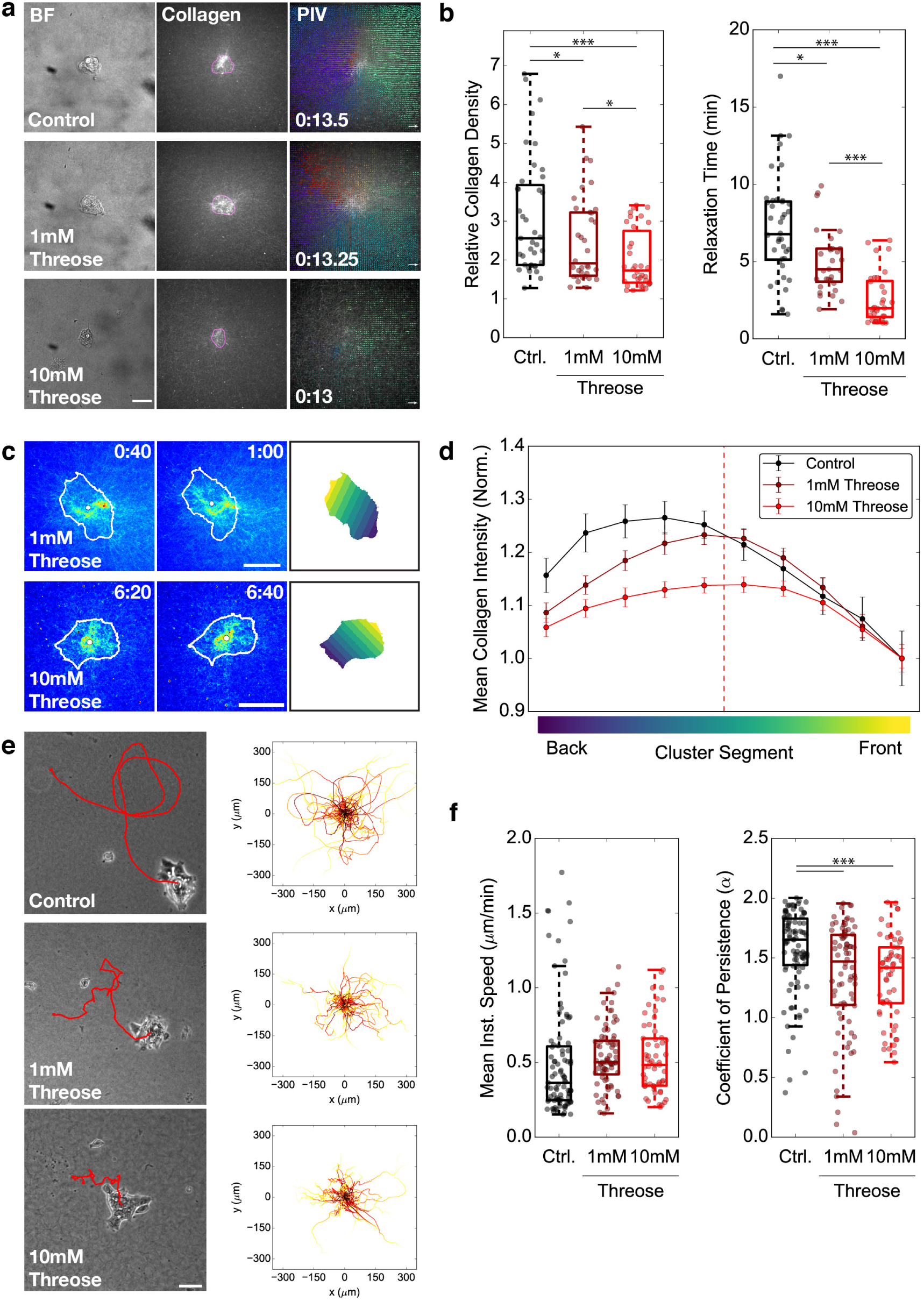
Collagen crosslinking decreases viscoelastic relaxation time and reduces persistent migration. **a.** Brightfield and confocal micrographs of cell clusters on fluorescent collagen networks with or without network crosslinking using threose and collagen images overlaid with PIV vectors following cell removal with Trypsin/NH_4_OH (see also Supplementary Movie 8-10). Magenta: segmented cell contour. Scale bar: 50*µ*m. HH:MM after addition of Trypsin/NH_4_OH. **b.** Boxplots of relative collagen density (*left*) and relaxation time (*τ*_*r*_, *right*) following cluster removal using Trypsin/NH_4_OH for conditions in *a*. Each dot represents one cluster that was rapidly removed. Data represents n = 38, 31, 32 clusters from N = 6, 4, 4 independent experiments. One-way ANOVA p<0.001, p<0.001. *p<0.05, ***p<0.001 for Tukey HSD post-hoc test. **c.** Sequential timepoints of cell clusters migrating on fluorescent collagen (brightest point projection) networks crosslinked with threose (see also Supplementary Movie 11,12). White line: segmented cluster contour. White dot: segmentation center of mass. Scale bar: 50*µ*m. HH:MM. *Right*: segmentation from the first time point shown, split into ten equal-length regions from the cluster front to back. **d.** Plot of the relative mean collagen intensity in different cluster segments during migration, for clusters migrating on collagen gels with or without pre-treatment of the crosslinker threose. Data represents mean±SEM averaged over all timepoints for n = 34, 30, 17 clusters from N = 5, 2, 3 independent experiments. Control data is the same as shown in Fig. 2e. **e.** Brightfield micrographs from cell cluster migration experiments on collagen networks, with the cell trajectory overlaid in red (see also Supplementary Movie 13-15). Images shown represent clusters at 16h after the start of the timelapse. *Right*: Overlaid migration trajectories for all clusters imaged and tracked, adjusted so that all trajectories start at the origin (0,0). **f.** Boxplots of mean instantaneous speed (*left*) and coefficient of persistence (*α*; *right*). Each dot represents one cluster trajectory. Data represents n = 91, 76, 57 clusters from N = 5, 2, 2 independent experiments. One-way ANOVA p>0.05, p<0.001. ***p<0.001 for Tukey HSD post-hoc test.

Based on our theoretical model, we expected that reducing the relaxation timescale would lead to a symmetric collagen density profile during cluster migration. Indeed, we found that the collagen density gradients on threose crosslinked gels were more symmetric than the controls (with a peak at the cluster center) and had lower magnitudes (Figure 4c,d, Supplementary Movies 11-12). These data support our model that reducing the substrate relaxation time leads to symmetric substrate deformations (Figure 3d).

Our model further predicts that migration should be less persistent on substrates with lower relaxation times. Consistent with this prediction, we observed that while clusters moved at similar speeds on control vs. threose-crosslinked collagen networks, clusters migrated significantly less persistently on crosslinked networks (Figure 4e,f; Supplementary Movies 13-15). Together, these data suggest that reducing the substrate relaxation time by crosslinking collagen networks leads to less persistent cluster migration, confirming one of the major predictions of our theoretical model.

### Single cells deform collagen networks less and migrate less persistently than clusters

Our theoretical model predicts that small clusters should not migrate persistently, due to the fact that smaller clusters exert less total force and are small compared to the substrate deformation length scale. As clusters appear to have characteristics of a large single cell (Figure S2), we compared cluster migration to single cell migration. To test whether single cells generate less stress on collagen networks, we performed 3D displacement microscopy for clusters and single cells (Figure 5a). We found that single cells exerted less total displacements on the substrate, indicating that they indeed exert less stress (Figure 5b). Previous work has suggested that the relaxation timescale of collagen networks depends on the amount of force exerted^19^. To test whether the lower stresses exerted by single cells could also affect collagen relaxation time, we rapidly removed single cells from collagen networks (Figure 5c, Supplementary Movie 16). We found that single cells generated lower initial collagen densities compared to clusters. Furthermore, collagen relaxation time following cell removal with Trypsin/NH_4_OH was lower for single cells compared to clusters (Figure 5d).

**Figure 5.**
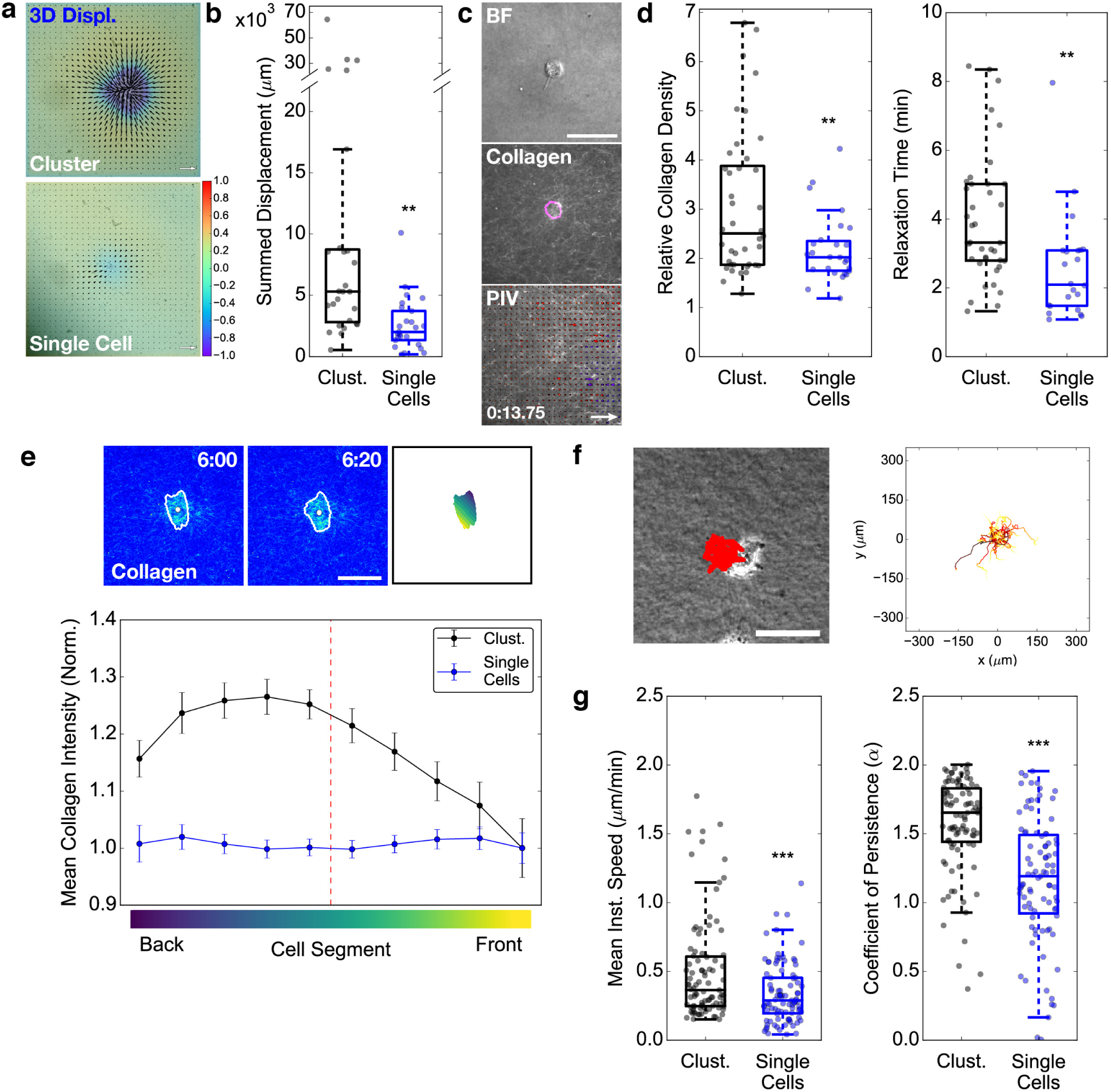
Single cells cannot sense local collagen gradients and migrate less persistently than clusters. **a.** 3D displacement microscopy of a cluster and single cell on PAA gels coated with collagen networks. Black arrows: *xy* displacements. Color scale: *z* displacements. **b.** Boxplot comparing summed 3D displacements for clusters vs. single cells. Each dot represents one cluster or single cell. Data represents n = 27, 25 clusters/cells from N = 4, 4 independent experiments. **p<0.01 for Welch’s t-test. **c.** Brightfield and confocal micrographs of a single cell on fluorescent collagen networks and collagen image overlaid with PIV vectors following cell removal with Trypsin/NH_4_OH (see also Supplementary Movie 16). Magenta: segmented cell contour. Scale bar: 50*µ*m. HH:MM. **d.** Boxplots of relative collagen density (*left*) and relaxation time (*τ*_*r*_, *right*) following removal of single cells from fluorescent collagen networks using Trypsin/NH_4_OH. Each dot represents one cluster or single cell that was rapidly removed. Data represents n = 38, 22 clusters from N = 6, 6 independent experiments. **p<0.01 for Welch’s t-test. Cluster data is the same as control data in Fig. 4b. **e.** *Upper panel*: Sequential timepoints of a single cell migrating on a fluorescent collagen network (brightest point projection), with segmentation split into ten equal-length regions from the cluster front to back (see also Supplementary Movie 17). White line: segmented cluster contour. White dot: segmentation center of mass. Scale bar: 50*µ*m. HH:MM. *Right*: segmentation from the first time point shown, split into ten equal-length regions from the cluster front to back. *Lower panel*: Plot of the relative mean collagen intensity in different cluster segments. Data represents mean±SEM averaged over all timepoints for 34,38 clusters from 5,5 independent experiments. Cluster data is the same as shown in Fig. 2e. **f.** Brightfield micrograph from single cell migration experiments on collagen networks, with the cell trajectory overlaid in red (see also Supplementary Movie 18). Image shown represents cell at 16h after the start of the timelapse. *Right*: Overlaid migration trajectories for all cells imaged and tracked, adjusted so that all trajectories start at the origin (0,0). **g.** Boxplots of mean instantaneous speed (*left*) and coefficient of persistence (*α*; *right*). Each dot represents one cluster or single cell trajectory. Data represents n = 91, 91 clusters/cells from N = 5, 3 independent experiments. ***p<0.001 for Welch’s t-test. Cluster data is the same as control data in Fig. 4f.

To determine whether single cells are able to generate collagen density gradients during migration, we performed live imaging of single cells migrating on fluorescent collagen gels (Figure 5e, *upper panel*; Supplementary Movie 17). We observed a flat collagen density profile across the length of single cells (Figure 5e, *lower panel*), suggesting that single cells are indeed too small to sense the collagen gradient during migration. Consistent with our model, we found that single cells migrated with lower mean instantaneous speeds and lower persistence compared to clusters (Figure 5f,g; Supplementary Movie 18). Together, these data support our model, which predicts that single cells, because they exert less traction stress and have a smaller area, migrate with less persistence compared to collectively migrating clusters.

## Conclusion

In this study, we identify a physical mechanism by which the material properties of a substrate can regulate migration dynamics of a cell cluster that otherwise lacks any intrinsic polarity. This mechanism relies intimately on the feedback between a cluster of cells and its substrate: the cell cluster induces a deformation of the substrate via traction forces, and the shape of the substrate deformation feeds back to control the direction of cell migration. The relatively slow viscoelastic relaxation of the substrate allows for persistent collective migration.

The role of substrate deformation during cell migration has been studied extensively using purely elastic PAA substrates^26–29^. A recent study comparing cell spreading and migration on elastic vs. viscoelastic PAA substrates found that some cell types can migrate with higher speed and more persistently on viscoelastic substrates^30^, consistent with our findings for cell cluster migration. Our study provides a mechanism for these findings, namely that increased relaxation time in viscoelastic substrates can lead to asymmetric substrate deformations during migration. This mechanism is similar in spirit to a model showing that apolar colloidal particles can become spontaneously self-propelled and swim in a highly persistent fashion as a result of changes in local solute concentrations due to hydrodynamic flows during particle motion^25^.

Recent work has shown that small groups of cells, similar to the clusters presented in this study, can collectively migrate up chemokine gradients^31, 32^. Collective chemotaxis is driven by polarized contractile myosin at the cluster rear^32^. In our study, persistent collective migration occurs spontaneously and in the absence of asymmetric myosin distributions (Figure S2c) and relies only on the physical interactions between the cluster and its substrate. It is likely, however, that the epithelial polarity mechanisms involved in keeping cell clusters together are required, as loss of these mechanisms lead to dissolution of the cell clusters into individual cells^13^.

Spontaneous persistent migration has recently been observed for cells migrating in uniform gradients of chemokine by self-generated chemotaxis^33, 34^. Because cells consume chemoattractant as they migrate, chemokine is depleted at the cell rear and remains highly concentrated at the cell front, effectively creating a sharp local chemokine gradient. This is similar to our mechanism in that local changes to the substrate/chemokine field during migration creates a polarity cue to continue migrating along the same direction. The finite diffusion of chemokine during self-generated chemotaxis is analogous to substrate relaxation time in our model.

Our theoretical model relies on a phenomenological coupling between the substrate deformation and the migrating cluster, which allows the cluster to actively respond to the substrate deformation. Although the precise identity of this coupling is not specified, there are a number of mechanisms that are likely to contribute. The high collagen density toward the rear of the cell could lead to increased adhesions due to higher ligand densities, which could result in more stable or numerous adhesions toward the cluster rear. Alternatively, because the minimum in collagen alignment is shifted toward the rear, this could favor protrusions formation along the aligned fibers and thereby promote persistent migration, as has been shown for exogenously aligned collagen networks^5^. These mechanisms are not mutually exclusive, and it is likely that several such factors contribute to the cluster’s response to asymmetric deformations in the collagen network.

Collective cell migration has emerged as a potential mechanism for tumor dissemination in early stages of metastasis. However, it is not clear whether collective migration offers any specific advantage over single cell migration during this early invasive stage. The results presented here provide a potential mechanism for increased migration efficiency for groups of cells migrating collectively vs. individual cells. This mechanism does not require specific acquisition of biochemical polarity for cell clusters, but simply relies on the physical interaction between groups of cells and stromalike networks of collagen-1. In summary, the experimental data and theoretical model presented here provides a simple physical mechanism for persistent collective cell migration on ECM networks, with potential impact in understanding stromal migration during early metastasis.

## Acknowledgements

The authors acknowledge the Cell and Tissue Imaging (PICT-IBiSA), Institut Curie, member of the French National Research Infractucture France-BioImaging (ANR10-INBS-04). We thank T. Kato and E. Sahai for the A431 MLC-GFP cell line, V. Marthiens and R. Basto for the Pericentrin antibody. We acknowledge M. Gómez-González and E. Latorre for developing the 3D PIV and TFM code. We thank H. Mohammadi, E. Sahai and all members of the Vignjevic lab for helpful discussions and comments on the manuscript. A.G.C. was supported by the European Moleclular Biology Organization (EMBO; ALTF 1582-2014 to A.G.C.) and Institut National du Cancer (INCa; PLBIO18-087 to D.M.V.). This project also received funding from the European Research Council (ERC) under the European Union Horizon 2020 research and innovation programme (grant agreement No 772487 to D.M.V.). A.M. and R.V. were funded by PHYMAX and POLCAM grants. X.T. was funded by Spanish Ministry for Science, Innovation and Universities MICCINN/FEDER (PGC2018-099645-B-I00, Severo Ochoa Award of Excellence), the Generalitat de Catalunya/CERCA program (SGR-2017-01602), Fundaci la Marat de TV3, and Obra Social “La Caixa”.

## Author Contributions

A.G.C., A.M., R.V. and D.M.V. designed the research and wrote the paper; A.G.C. carried out most of the experiments and image analysis; A.M. and R.V. created the theoretical model; C.J. performed and analyzed the migration and 2D TFM experiments; A.S. performed preliminary experiments; C.P.G. and X.T. provided reagents, software and technical assistance for the TFM and 3D displacement experiments; all authors discussed the results and manuscript.

