## Supplementary Information for "Viscoelastic relaxation of collagen networks provides a self-generated directional cue during collective migration"

### Methods

**Cell culture and preparation.** A431 cells were cultured in DMEM (Gibco/Life Technologies) supplemented with 10% fetal bovine serum (FBS; Gibco). A431 wild type cells were a gift from C. Rosse and P. Chavrier (Institut Curie) and were authenticated using the Gene Print 10 System (Promega). A431 MLC-GFP cells were a gift from T. Kato and E. Sahai (Francis Crick Institute); parental A431 lines were authenticated by STR profiling. Cells were passaged 2-3 times per week and tested for mycoplasma every 2 weeks.

Cell clusters were prepared by adding $\sim$2x10$^6$ cells to a 10cm dish that was pre-coated with 1% Agarose in PBS and filled with DMEM + 10% FBS + 2x Anti/Anti (ThermoFisher). Cells were cultured in these non-adherent conditions overnight prior to being seeded on substrates for imaging. Before seeding, large clusters and single cells were removed: (1) the medium containing the clusters from the agarose-coated dish was pipetted into a 15mL falcon tube, (2) large clusters were allowed to sediment for 1min before all but 0.5mL of the medium was transferred to a new 15mL falcon tube, (3) cells were centrifuged briefly at $\sim$140xg and the supernatant was discarded, (4) the pellet was resuspended in 0.5mL medium; $\sim$15$\mu$l cluster solution was used for seeding. For single cell experiments, cells were detached, and 2x10$^3$ cells were seeded on collagen networks. Cells were seeded in AB+ medium (DMEM + 10%FBS + 2x Anti/Anti + 0.125% Metronidazole (w/v) + 4$\mu$g/ml Ciprofloxacin) and imaged 2-3 days after seeding.

**Preparation of collagen and poly-A-acrylamide substrates.** For experiments with cells/clusters on collagen networks, 200-600$\mu$l of neutralized rat tail collagen-1 solution (2mg/ml; Corning) was pipetted onto 3.5cm glass bottom dishes (FluoroDish, WPI) and incubated at 37°C with humidity for 20min before adding 2mL AB+ medium with cells/clusters. Glass bottom dishes were pre-coated with 3-aminopropyltrimethoxysilane (diluted 1:2 with water, Sigma) and glutaraldehyde (0.5% in PBS) and washed well with water.

For experiments using coated PAA gels, PAA solution for 0.5kPa gels were prepared by mixing 50$\mu$l 40% acrylamide (Bio-Rad), 7.5$\mu$l 2% bis-acrylamide (Bio-Rad), 2.5$\mu$l 10% ammonium peroxodisulfate (VWR) and 0.25$\mu$l tetramethylethylenediamine (Euromedex) in PBS (full volume 500$\mu$l). Then, 16$\mu$l of solution was pipetted onto silane/gluteraldehye-coated glass bottom dishes and covered with a round 18mm diameter coverslip. Gels were incubated for 60min before washing with PBS and removing the coverslip. For TFM experiments, 10$\mu$l 2% 0.5$\mu$m diameter green fluorescent polystyrene beads (Invitrogen/ThermoFisher) were also added.

PAA gels were coated by incubating the gel with 2mg/ml sulfo-SANPAH (Sigma-Aldrich/Merck) under ultraviolet light (365nm; 10cm from source) for 10min. The gel was then washed 2x3min with 10mM HEPES and 1x3min with PBS to remove the excess sulfo-SANPAH. For coating with thin collagen networks, 100$\mu$l of neutralized 2mg/ml collagen solution was added to cover the PAA gel and all excess collagen solution was immediately removed. Collagen was allowed to polymerize at 37°C with humidity for 20min. For coating with monomeric collagen, the desired concentration of collagen (100$\mu$g/ml for Fig. 1, different concentrations for Fig. S1) was diluted in 0.2% Acetic Acid and pipetted to cover the PAA gel. The dish was then incubated at 37°C with humidity for 1hr and washed briefly with PBS before adding AB+ medium and clusters.

**Immunostaining.** For immunostaining of Rac1 in cell clusters, dishes containing clusters on collagen networks were washed briefly with phosphate buffered saline (PBS) and then simultaneously fixed and extracted using a solution of 4% PFA, 5% Sucrose and 0.3% Triton X-100 in PBS for 5min at room temperature. Cells were then further fixed with 4% PFA and 5% Sucrose in PBS for 40min at room temperature before washing 2x5min with PBS. Cells were stained with a primary antibody solution of 1:100 Mouse-anti-Rac1 antibody (610650, BD Biosciences) in PBS overnight. The following day, cells were washed 5x30min with PBS-T (PBS + 0.1% Tween20) and then stained with a secondary antibody solution of 1:200 Goat-anti-MouseIgG-Alexa568 (ThermoFisher) + 1:200 DAPI + 1:200 Phalloidin-Alexa633 (ThermoFisher) in PBS overnight. Samples were then washed 5x30min with PBS-T and incubated in PBS + 2xAnti/Anti until imaging. Rac1-stained clusters were imaged in 3D using an upright spinning disc confocal (Zeiss) with a 63x dipping lens (NA 1.0) and 1$\mu$m z-steps.

For immunostaining of centrosomes, cell clusters were washed briefly and fixed with 4% PFA in PBS for 40min at room temperature. Samples were then washed briefly with PBS, extracted with 0.5% Triton X-100 in PBS for 5min and then washed 3x5min in PBS. Cells were stained with a primary antibody solution of 1:500 Rabbit-anti-Pericentrin antibody (ab4448, Abcam; gift from V. Marthiens and R. Basto) in PBS overnight. The following day, cells were washed 5x30min with PBS-T and then stained with a secondary antibody solution of 1:200 Goat-anti-RabbitIgG-Alexa568 (ThermoFisher) + 1:200 DAPI + 1:200 Phalloidin-Alexa633 (ThermoFisher) in PBS overnight. Samples were then washed 5x30min with PBS-T and incubated in PBS + 2xAnti/Anti until imaging. Pericentrin-stained clusters were imaged in 3D using an inverted Eclipse Ti-E microscope (Nikon) with Spinning disk CSU-W1 (Yokogawa) integrated in Metamorph software by Gataca Systems with a 40xW immersion lens (NA 1.15) and 1$\mu$m z-steps.

**Imaging and analysis of migration experiments.** Two to three days after seeding cells/clusters, samples were imaged using an Inverted Eclipse Ti-E microscope (Nikon) with a motorized stage and a 10x (NA 0.3) objective for $\sim$16h. Multiple cells/clusters were imaged using the Multi-Dimensional Imaging module in Metamorph. To obtain migration trajectories, cells/clusters in brightfield timelapse images were segmented and tracked using Ilastik[35]. From the tracked segmentations, the trajectories were determined from the center of mass of the segmentation at each time frame. Cluster fusion or fission events were not included in the analysis, and only trajectories before or after these events were considered. Analysis and plotting of the trajectories were performed using custom software written in Python.

**Imaging and analysis of cortical myosin intensity during migration.** A431 MLC-GFP clusters were imaged using an inverted Eclipse Ti-E microscope (Nikon) with Spinning disk CSU-W1 (Yokogawa) integrated in Metamorph software by Gataca Systems with a 40xW immersion lens (NA 1.15) and 1$\mu$m z-steps for $\sim$16h. Multiple clusters were imaged in 3D using the Multi-Dimensional Imaging module in Metamorph. To determine cortical intensities of MLC-GFP, we first registered the 3D timelapse images in the z-axis using a custom macro in FIJI. We then segmented cluster timelapse images automatically using custom software written in Python. Briefly, each image was blurred using a Gaussian filter and thresholded using Otsu's method. The segmentations were further refined using binary morphology operations to remove noise and to smooth edges. The cells were then tracked frame-to-frame to extract the trajectory using the center of mass for each segmented image. Using the segmentation, we defined the cortex as the region around the cluster periphery from the outer boundary of the cluster to a contour 5$\mu$m inside of the boundary. For each cortex pixel, we determined the angle of that pixel with respect to the cluster center of mass. The cortex intensity around the cluster was then determined by finding the maximum cortex intensity value within angular bins of 10°. This cortical intensity was normalized to the average intensity in the region of the cluster inside of the cortex. For averaging the angular cortex intensities with respect to the angle of migration, we subtracted the trajectory angle from the angle value for each cortex pixel and determined the binned cortex intensities as described above.

**Analysis of centrosome position in cell clusters.** Relative centrosome positions in cell clusters were determined by segmenting the cluster volume (Phalloidin channel), nuclei (DAPI channel) and centrosomes (Pericentrin channel) in 3D. Initially a segmentation probability map was generated for each channel using Ilastik. Each channel was segmented by thresholding the probability map and performing binary morphology operations. Each nucleus was then paired with the closest centrosome (typically adjacent to the surface of the nucleus). Centrosome orientation was defined by a unit vector from the nucleus center of mass to the centrosome center of mass for each nucleus-centrosome pair.

**Imaging and analysis of fluorescent collagen networks during migration.** Monomeric collagen-1 was labeled with TAMRA as previously described[36]. For fluorescently-labeled collagen networks, TAMRA collagen was mixed with unlabeled collagen at a ratio of 1:5 and neutralized;

200$\mu$l of collagen solution was pipetted onto silane/glutaraldehyde-coated glass bottom dishes and polymerized as described above. Single cells or clusters were then seeded. After 2-3 days, the cells were labeled with CellTracker Green (ThermoFisher) at 1:2000 in serum-free DMEM for 30min at 37°C with 5% $CO_2$ and humidity. Following incubation, the medium was replaced with fresh AB+ medium. Samples were imaged $\sim$1hr later by 3D spinning disc microscopy using an inverted Eclipse Ti-E microscope (Nikon) with Spinning disk CSU-W1 (Yokogawa) integrated in Metamorph software by Gataca Systems with a 40xW immersion lens (NA 1.15) and 1$\mu$m z-steps at 20min time intervals for $\sim$16h. Multiple cells/clusters were imaged in 3D using the Multi-Dimensional Imaging module in Metamorph.

Segmentation and tracking were performed on brightest point projections of CellTracker images using a custom algorithm in Python similar to those used for A431 MLC-GFP images. To determine the angular difference between clusters and the collagen center of mass, the angle from the cluster segmentation center of mass to the weighted center of mass of the collagen intensity within the segmentation region was determined. The difference between this angle and the trajectory angle was taken to be the angular difference.

To measure the local nematic order outside of the cell region (Figure S5), collagen fiber orientations were determined for a single $z$-slice at the surface of the collagen network using CT-FIRE[37]. Next, the image frame was split into a grid of square boxes, each with a side length of 120px (33$\mu$m). Within each box, we calculated the nematic order using a nematic director $\mathbf{n}$ defined by the mean filament orientation in that region. We calculated the nematic order scalar parameter considering all fibers contained within the box and not intersecting the cluster segmentation as $S(x,t) = \frac{1}{2}\langle 3\cos^2\theta_m - 1\rangle$, where $\theta_m$ is the angular difference between each fiber orientation and the nematic director $\mathbf{n}$.

To one-dimensionalize the collagen density signature, cell segmentations at each time point were divided into ten regions of equal length from the front to the rear of the cluster, according to the angle of migration at that time step. Regions were determined by making a bounding box around the segmentation, rotated with the trajectory angle, and separating this bounding box into ten equal rectangles. The segmentation pixels intersecting these rectangles were used in determining the mean collagen intensity at each region. The nematic order for each region was determined for all fiber orientations in that region (obtained using CT-FIRE) using the migration trajectory angle as the nematic director $\mathbf{n}$. The nematic order was only calculated for regions containing $>3$ filament orientations. For each time point, the collagen density and nematic order were normalized to the mean of the ten regions. The collagen density peak was determined by fitting the averaged collagen density data with a Gaussian function and extracting the center position, $\mu$. The nematic order trough was determined by fitting an inverted Gaussian function. The reported peak offset for collagen density and nematic order was taken as the relative peak position multiplied by the mean segment length.

**2D traction force microscopy.** For 2D traction force experiments, cell clusters were seeded on 0.5kPa PAA gels containing fluorescent beads and coated with a thin collagen-1 network as described above. Clusters and beads were imaged by brightfield and epifluorescence microscopy, respectively, overnight using an Inverted Eclipse Ti-E microscope (Nikon) with a full motorized stage and a 10x (NA 0.3) objective. Multiple cells/clusters were imaged using the Multi-Dimensional Imaging module in Metamorph. A single $z$-slice was acquired every 15min for multiple stage positions and imaged overnight for $\sim$16h. The following day, the cells were removed by briefly rinsing with PBS and incubating for 20min in 0.5x TrypLE Express (ThermoFisher) and 1N $NH_4OH$. The dish was then briefly rinsed with PBS and the collagen-1 network was removed by incubating for 20min in a solution of $\sim 50\mu$g/ml Collagenase type 3 (Worthington Biochemical) in PBS. All incubations to remove cells and the collagen network were performed directly on the microscope, being careful not to disturb the position of the imaging dish. An additional image was then taken to serve as a reference image for measuring bead displacements. Beads displacements were extracted by PIV and tractions were calculated from displacement fields by Fourier-transform, assuming finite gel thickness using custom software in Matlab[38].

Clusters were segmented and tracked over time from brightfield images using Ilastik[35]. The migration trajectories were pooled with the cluster migration data described above. Traction force linescans were extracted by cubic spline interpolation of the traction field. The linescans ran from each contour pixel to the segmentation center of mass and extended outward $40\mu$m beyond the segmentation boundary. For averaging around a single cell, linescans were aligned to the countour boundary of each linescan. The peak traction magnitudes was taken as the highest traction value in the linescan between $20\mu$m outside of the cluster boundary and $10\mu$m inside the cluster boundary. To quantify the peak traction magnitude around the cluster according to the angle, each contour pixel was assigned an angle according to the vector pointing from the center of mass of the segmentation to the contour pixel. To average over time and many clusters, the migration trajectory angle was subtracted from each contour angle (as for the MLC-GFP analysis above), and the traction peak was averaged along binned angles to achieve a mean polar plot of the peak traction magnitude with respect to migration direction. To calculate the front-to-back ratio of peak traction magnitudes, the mean peak traction from the front quadrant (315°-45°) was divided by the mean of the peak tractions from the rear quadrant (135°-225°).

For calculating the tractions along the axis of migration, each traction vector was projected along the migration axis, and the magnitude of this projected vector was plotted. Linescans of the projected tractions were generated by making a bounding box of 500px x 500px ($\sim$320$\mu$m x 320$\mu$m) to fit around the segmented cluster. The bounding box was then split into 500 regions from the front to the back, and the tractions (expanded to the image grid by nearest neighbor interpolation) within each segment were averaged (mean) to determine the traction value at each point along the cluster length. The raw values were smoothed using a moving window with size = 21px. To include the linescan edges in the smoothed data, the linescan was padded at the start and end using the start and end values, respectively. The linescan was then cropped to start and end when the data reached 0.

**3D deformation microscopy.** 3D deformation microscopy experiments were set up in the same way as for 2D traction force experiments. Single cells or cell clusters were seeded on 0.5kPa PAA gels containing fluorescent beads and coated with a thin collagen-1 network. Samples were imaged by 3D spinning disc microscopy using an Inverted Eclipse Ti-E microscope (Nikon) with Spinning disk CSU-W1 (Yokogawa) integrated in Metamorph software by Gataca Systems with a 40xW immersion lens (NA 1.15). A single 3D stack with $1\mu$m z-steps was collected. Then the cells and collagen network were detached as described above, and a reference image stack was collected. Bead displacements along the $x$,$y$ and $z$ axes were extracted by a custom 3D PIV algorithm in Matlab. The summed displacement was the sum of the magnitude of all $xyz$ vectors in the imaging frame.

**Collagen and stress relaxation following cell removal.** To determine the collagen relaxation timescale, cells or clusters were seeded on TAMRA-collagen networks as described above. A single 3D image stack was recorded by spinning disc microscopy for each cell or cluster, with 35 $2\mu$m z-steps. Cells were then removed by treatment with 0.5x TrypLE Express and 1N $NH_4OH$. Imaging was resumed as soon as possible after the sample was stable (5-10min), noting precisely the time between cell removal and restarting imaging. Cell or cluster regions were segmented manually based on the first (pre-Trypsin/$NH_4OH$) image. The collagen density at each time point was taken as the mean intensity of a brightest point projection of the collagen intensity within the segmentation region normalized to a reference region with the same shape as the segmentation. PIV analysis of collagen relaxation for sequential frames was performed using the openpiv library in Python. To determine the relaxation of stresses on the substrate following cell removal, 3D deformation microscopy was performed as described above. A single 3D image stack of clusters or single cells and their underlying collagen network/PAA gel was recorded by spinning disc microscopy as described above. Cells were then removed using Trypsin/$NH_4OH$ and imaging was resumed after 5-10min. The relaxation timescale $\tau_r$ was extracted by fitting the displacement/density/PIV data over time with the exponential decay function $I = A\exp(-t/\tau) - C$.

**Preprocessing, image analysis and visualization.** Preprocessing of image stacks was performed using custom macros in FIJI/ImageJ[39]. For image and data analysis and visualization in Python, the following packages were used: numpy, scipy, pandas, scikit-image, opencv, astropy, shapely and matplotlib. For boxplots, dots represent individual measurements as described in the figure legends. Boxes represent the interquartile range (IQR; Q1 to Q3) and the line represents the median. Whiskers extend to the furthest point within 1.5*IQR away from Q1 or Q3.

**Data availability** The authors declare that all data supporting the findings of this study are available within the paper and its supplementary information files and from the corresponding authors upon reasonable request.

**Code availability** Custom software used to analyze images and data will be made available upon reasonable request.

### Supplementary Note: details on the theoretical model

#### Introduction and summary of the results

We describe here a simple model of cell cluster migration on a viscoelastic substrate, which shows that an apolar cell cluster can perform persistent self propelled motion in absence of any internal self polarization mechanism, and in absence of any external polar cue. The model is general and does not rely on a specific cell type; its aim is to identify the minimal ingredients required for persistent polarized migration of an apolar cell cluster. As we explain below, the main ingredient relies on the active coupling of the apolar cluster to the apolar viscoelastic medium, which we show is sufficient to induce a symmetric breaking in the coupled cluster/substrate system, and thereby persistent motion of the cluster.

In its simplest form, the model first describes the cell cluster as an isotropic active particle, whose position of the center of mass along a reference axis is denoted by $x_c$. The cluster actively interacts with the underlying viscoelastic substrate: we assume that the cluster, by exerting an active stress on the substrate of intensity prametrized by $f$, is the source of a scalar structural perturbation $S(x,t)$ in the substrate; without being specific, for a contractile cluster seeded on an ECM-like fibrilious polymeric substrate, $S(x,t)$ can be a measure of the scalar nematic order parameter in the substrate, and/or a local variation in density. This perturbation is described phenomenologically in the model and characterized by a relaxation time $\tau_r$ and a localization length scale $\ell$; this description is based on a minimal description of the substrate rheology as a linear elastic solid with viscous relaxation. In turn, we assume that the cluster actively responds to the substrate perturbation $S(x,t)$. We do not aim at identifying the microscopic cellular mechanisms involved in this mecanosensitive response, which could be a response to stiffness (i.e. indirectly density) and/or order in the substrate, as was reported at the single cell and cell assemblies levels in various cell types [28,41,42]. Introducing a phenomenological coupling $\zeta$, which parametrizes the active response of the cluster to the perturbation, we write such response in its simplest linear form $v_c = \dot{x}_c = \zeta \partial_x S|_{x_c} + \eta(t)$, where fluctuations can be encoded in a noise term $\eta$.

The main results of this model can be summarized as follows : (i) in the absence of noise, a (supercritical) instability occurs for a critical value $\zeta_c$ of the activity parameter (with all other parameters held fixed): for $\zeta < \zeta_c$, one has $v_c = 0$, and the cluster cannot migrate persistently; for $\zeta > \zeta_c$, one has $v_c \propto \pm(\zeta - \zeta_c)^{1/2}$. Importantly, we show that the instability threshold scales as $\zeta_c \sim (f\tau_r)^{-2}$ and is therefore critically controlled by the relaxation time of the gel – at a given level of activity, persistent motion can be enhanced simply by engineering a gel with a $\tau_r$ larger than the critical relaxation time $\tau_c$. The dependence on the cluster size $L$ can also be inferred by the model, by noting that both $\zeta$ and $f$ are increasing functions of $L$ : persistent motion is therefore predicted to occur only for large enough clusters. (ii) In the presence of noise, the model therefore predicts a Brownian-like random migration with negligible persistence time for $\zeta < \zeta_c$ (or $\tau < \tau_c$), in turn, for $\zeta > \zeta_c$ (or $\tau_r > \tau_c$), it predicts a persistent motion, whose persistence time increases with $\zeta - \zeta_c$, and therefore with $\tau_r$. These results are qualitatively in agreement with experimental observations. A qualitative picture can be useful: by contracting, the cluster effectively induces a local bumpy perturbation in the substrate; in turn, the cluster can slide downhill along the perturbation profile. For a long enough relaxation time of the substrate, it is found that the cluster can surf the bump it induces at constant speed.

Last, we provide a general theoretical framework that describes the active dynamics of an apolar system localised in space (the cluster), that we parametrize by a phase field $\phi(x, t)$, which interacts with a generic viscoelastic nematic substrate. This description is fully general, and is shown in particular to encompass the simplest version of the model; its analysis shows that the mechanism leading to persistent motion can be generalized and could be at work in other active systems, living or artificial. Furthermore, this more general model also suggests that larger clusters are more persistent and lead to greater substrate anisotropy, consistent with the experiment.

**Minimal model of apolar active particle coupled to a viscoelastic environment**

In this section, we write down a simple, one-dimensional heuristic model in which we describe the cell cluster by the position of its centroid $x_c$. The cell cluster induces a structural perturbation in the substrate, which we describe by a non-dimensional scalar $S(x, t)$:

$$\partial_t S = -\frac{1}{\tau_r}[S - \ell^2 \partial_x^2 S] + f\delta(x - x_c). \tag{2}$$

Here, the last term on the R.H.S. is the forcing due to the cluster with a strength $f$ which has the dimension of a speed. This equation is derived below in the case where $S$ is the scalar nematic order parameter, or local variation of gel density. The active response of the cluster to the perturbation is written phenomenologically as

$$\dot{x}_c = -\zeta \partial_x S|_{x_c}. \tag{3}$$

We now consider a cluster that moves steadily at speed $v_c$. Defining $\mu = x - v_c t$, and noting that $x_c = v_c t$, we obtain a pair of self-consistent equations for determining $v_c$:

$$\left[\frac{1}{\ell^2} - \frac{v_c \tau_r}{\ell^2}\partial_\mu - \partial_\mu^2\right] S = f\frac{\tau}{\ell^2}\delta(\mu) \tag{4}$$

$$v_c = -\zeta \partial_\xi S|_0 \tag{5}$$

From (4), and assuming that $S$ vanishes at large scales, one obtains

$$S(\mu) = -e^{-\frac{\mu(v_c\tau_r + \sqrt{4\ell^2 + v_c^2\tau_r^2})}{2\ell^2}} \frac{f\tau_r}{\sqrt{4\ell^2 + v_c^2\tau_r^2}} \left(e^{-\frac{\mu\sqrt{4\ell^2+v_c^2\tau_r^2}}{\ell^2}}\Theta[-\mu] + \Theta[\mu]\right), \tag{6}$$

which, when inserted into (5) yields, in addition to $v_c = 0$, the solutions

$$v_c = \pm\frac{\sqrt{f^2\zeta^2\tau_r^4 - 16\ell^6}}{2\ell^2\tau_r}. \tag{7}$$

This has the form

$$v_c = \pm\frac{f\tau_r}{2\ell^2}\sqrt{\zeta^2 - \zeta_c^2} \tag{8}$$

where $\zeta_c = 4\ell^3/f\tau_r^2$. That is, as a function of the activity parameter, $v_c$ undergoes a supercritical pitchfork bifurcation at $\zeta_c$ and assumes a non-zero value. Eq. (8) implies that when $\zeta \gg \zeta_c$, $v_c \sim \zeta$. When all other quantities are held constant, $v_c \sim \tau_r$ at large $\tau_r$ and $v_c \sim 1/\ell^2$.

So far we assumed that the forcing of the polymeric substrate due to the cell cluster is localised purely at the centroid of the cluster. We now relax this hypothesis and demonstrate that the basic mechanism leading to the instability remains correct even when the source of the perturbation $\delta(x - x_c)$ in (2) is not localized, but replaced by a general function $g(x - x_c)$ peaked at $x_c$ and vanishing at large scales, and with its integral normalized to 1. That is,

$$\partial_t S = -\frac{1}{\tau_r}[S - \ell^2\partial_x^2 S] + fg(x - x_c). \tag{9}$$

Importantly, $g$ is assumed to be even, meaning that the cluster has no intrinsic polarity. We assume that $\int_{-\infty}^{\infty} g(y)dy = 1$. To reduce the number of parameters, we now rescale space by $\ell$ i.e., define $\tilde{x} = x/\ell$ and time by $\tau_r$, i.e. $\tilde{t} = t/\tau_r$. Further, defining $\xi = \tilde{x} - \bar{v}_c\tilde{t}$, where $\bar{v}_c = v_c\tau/\ell$, we get the equations of motion

$$\left[-1 + \bar{v}_c\partial_\xi + \partial_\xi^2\right] S = -\tilde{f}g(\xi) \tag{10}$$

where $\tilde{f}$ is the non-dimensionalised version of $f$ and depends linearly on $\tau_r$, $g(\xi)$ is a non-dimensionalised version of the function $g(x - x_c)$ and

$$\bar{v}_c = -\tilde{\zeta}\partial_\xi S|_0 \tag{11}$$

where $\tilde{\zeta} = \zeta\tau_r/\ell^2$. We can now solve (10) using the Green's function

$$\mathcal{G}(\xi, \xi') = -\frac{1}{\sqrt{\bar{v}_c^2 + 4}} e^{\frac{\xi-\xi'}{2}(-\bar{v}_c+\sqrt{\bar{v}_c^2+4})} \left[e^{(\xi'-\xi)\sqrt{\bar{v}_c^2+4}}\Theta(\xi - \xi') + \Theta(\xi' - \xi)\right]. \tag{12}$$

with $S(\xi) = \tilde{f}\int_{\infty}^{\infty} \mathcal{G}(\xi, \xi')g(\xi')$. The speed can then be calculated as

$$\bar{v}_c = \tilde{\zeta}\partial_\xi S|_0 = \tilde{\zeta}\partial_\xi \int_{-\infty}^{\infty} \mathcal{G}(\xi, \xi')\tilde{f}g(\xi')d\xi'|_0 = \tilde{\zeta}\int_{-\infty}^{\infty} \partial_\xi[\mathcal{G}(\xi, \xi')]\tilde{f}g(\xi')d\xi'|_0. \tag{13}$$

Here,

$$\partial_\xi[\mathcal{G}(\xi, \xi')] = -\frac{1}{\sqrt{4 + \bar{v}_c^2}} e^{\frac{-\bar{v}_c+\sqrt{4+\bar{v}_c^2}}{2}(\xi-\xi')}$$
$$\left[\left\{e^{\sqrt{4+\bar{v}_c^2}(\xi'-\xi)} - 1\right\}\delta(\xi - \xi') - \frac{\bar{v}_c + \sqrt{4 + \bar{v}_c^2}}{2} e^{\sqrt{4+\bar{v}_c^2}(\xi'-\xi)}\Theta(\xi - \xi') + \frac{-\bar{v}_c + \sqrt{4 + \bar{v}_c^2}}{2}\Theta(\xi' - \xi)\right]. \tag{14}$$

Using this, one obtains

$$\bar{v}_c = \tilde{\zeta}\int_{-\infty}^{\infty} \frac{e^{\frac{\bar{v}_c-\sqrt{4+\bar{v}_c^2}}{2}\xi'}\left[(\bar{v}_c + \sqrt{4 + \bar{v}_c^2})e^{\sqrt{4+\bar{v}_c^2}\xi'}\Theta(-\xi') + (\bar{v}_c - \sqrt{4 + \bar{v}_c^2})\Theta(\xi')\right]}{2\sqrt{4 + \bar{v}_c^2}}\tilde{f}g(\xi')d\xi'. \tag{15}$$

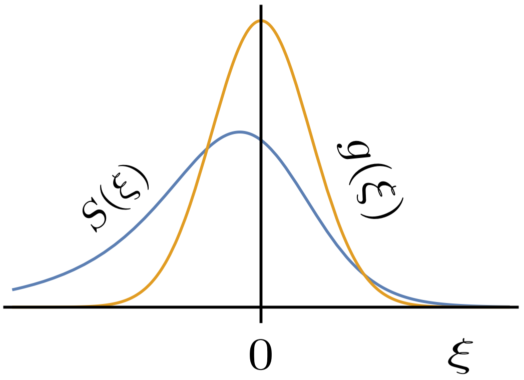


Figure T1: The cluster induces a symmetric contractile stimulus $g$ localized at the cluster centre. Because of the finite relaxation time $\tau_r$, the response of the substrate (perturbation $S$) is delayed. For a moving cluster, this delay causes a space shift and asymmetry of the perturbation $S$.

Since the kernel in (15) is not symmetric under $\xi' \to -\xi'$, a function $g(\xi')$ which is symmetric and has a maximum at $\xi' = 0$ should yield a non-zero value of the integral. See Fig. T1 for an example. Since we demand that $g(\xi')$ has a maximum at $\xi' = 0$, $g(\xi')$ must remain (at least) finite as $|\xi'| \to \infty$. The value of the integral is finite for all such $\tilde{f}(\xi')$. Furthermore, since the integral goes to $0$ as $|\bar{v}_c| \to \infty$, a nontrivial (non-zero) solution of the self-consistent equation for $\bar{v}_c$, (15) *must* exist when the slope of the R.H.S. of (15) with respect to $\bar{v}_c$ exceeds $1$ at $\bar{v}_c = 0$. That is, the cluster spontaneously transitions to a symmetry-broken steadily-moving state beyond a critical activity $\tilde{\zeta}_c$ given by

$$\frac{4}{\tilde{\zeta}_c} = \int_{-\infty}^{\infty} \tilde{f} g(\xi') \left[ e^{\xi'}(1+\xi')\Theta(-\xi') + e^{-\xi'}(1-\xi')\Theta(\xi) \right] d\xi' \tag{16}$$

Notice that the integral on the R.H.S. does not diverge for any reasonable function which has unique maximum at $\xi' = 0$. (Notice that the term in the square bracket vanishes as $|\xi'|e^{-|\xi'|}$ as $|\xi'| \to \infty$.) This equation therefore always has a solution – again implying that there is always

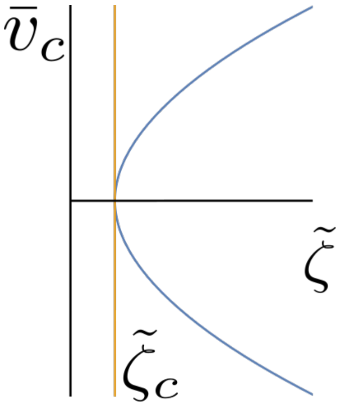


Figure T2: $\bar{v}_c$ undergoes a supercritical pitchfork bifurcation at $\tilde{\zeta} = \tilde{\zeta}_c$. Here, $\bar{v}_c$ is plotted as a function of $\tilde{\zeta}$ for $g(\xi') = e^{-\xi'^2/2}/\sqrt{2\pi}$.

a critical (non-zero) $\tilde{\zeta}_c$ beyond which the cell cluster starts moving, *irrespective* of the precise form of the forcing function $g(\xi)$. We can also infer the scaling of $\tilde{\zeta}_c$ on $\tau_r$ directly from this: $\tilde{f} \sim .f\tau_r$ which implies $\tilde{\zeta}_c \sim (f\tau_r)^{-1}$ and using the definition of $\tilde{\zeta}$, $\zeta_c \sim (f\tau_r)^{-2}$ as given in the main text. We demonstrate the supercritical bifurcation diagram implied by (15) for a profile with $g(\xi) = e^{-\xi^2/2}/\sqrt{2\pi}$ with $\tilde{f} = 1$ in Fig. T2.

It is difficult to directly read-off the exact dependence of the speed of the cluster on $\tau_r$ and $\ell$ as it depends on the form of the forcing function. To show this explicitly, we revert to the dimensional variables and write the fully dimension restored version of (15):

$$v_c = \frac{\zeta f \tau_r}{\ell^2} \int_{-\infty}^{\infty} d\mu' g(\mu') \frac{e^{-\frac{\mu'\left(-v_c\tau_r + \sqrt{4\ell^2 + v_c^2\tau_r^2}\right)}{2\ell^2}}}{2\sqrt{4\ell^2 + v_c^2\tau_r^2}} \left[ e^{\frac{\mu'\sqrt{4\ell^2+v_c^2\tau_r^2}}{\ell^2}} \left(v_c\tau_r + \sqrt{4\ell^2 + v_c^2\tau_r^2}\right)\Theta(-\mu') + \left(v_c\tau_r - \sqrt{4\ell^2 + v_c^2\tau_r^2}\right)\Theta(\mu')\right] \quad (17)$$

where $\mu = x - v_c t$ as defined earlier. It is clear that the detailed dependence of the solution of this equation on $\tau_r$ is complicated, in general. We now specialise to a Gaussian profile for

$g(\mu) \equiv e^{-\mu^2/2\sigma^2}/\sqrt{2\pi}\sigma$ such that we recover the results we obtained for the $\delta$ function forcing when $\sigma \to 0$. In this case, the condition for the cluster motility becomes

$$\frac{f\zeta\tau_r^2}{2\sqrt{2\pi}\ell^5}\left[-2\ell\sigma + e^{\frac{\sigma^2}{2\ell^2}}\sqrt{2\pi}(\ell^2+\sigma^2)\mathrm{erfc}\left(\frac{\sigma}{\sqrt{2}\ell}\right)\right] > 1. \tag{18}$$

This clearly implies that increasing the relaxation time $\tau_r$ or the activity $\zeta$ promotes the motility of the cluster and the critical activity $\zeta_c \sim \tau_r^2$. In other words, if we increase $\tau_r$ without modifying any other parameter, at a finite $\tau_r = \tau_c$ a $v_c \neq 0$ solution sets in and the particle starts moving spontaneously. We have checked that this conclusion is unmodified even for more general forcing functions.

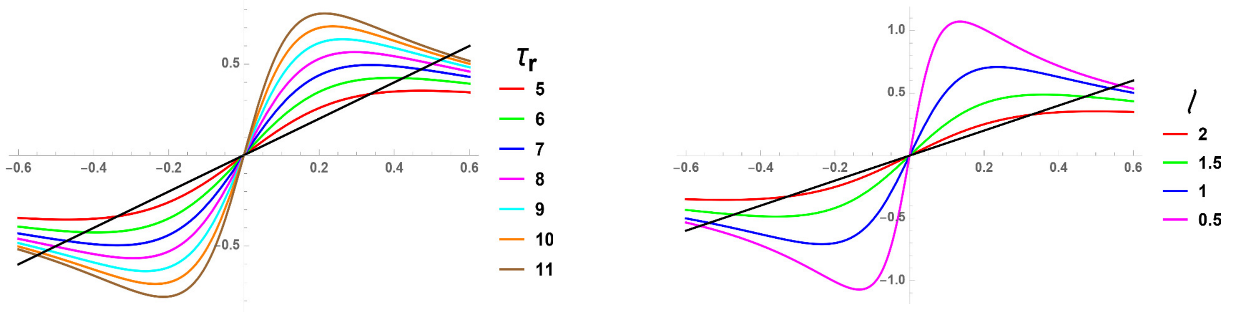


(a) Solutions of (17) with $\ell = 1$, $\zeta = 1$, $f = 1$, and $\sigma = 1$

(b) Solutions of (17) with $\tau_r = 10$, $\zeta = 1$, $f = 1$, and $\sigma = 1$

Figure T3: The solutions of (17) for multiple $\ell$ and $\tau_r$; the intersection of the black straight line with the curves for different $\tau_r$ is the solution of $v_c$ for that $\tau_r$. This demonstrates that $v_c$ increases with $\tau_r$ before asymptotically going to a constant value. Similarly, it decreases with $\ell$ and asymptotically goes to a constant value for large $\ell$.

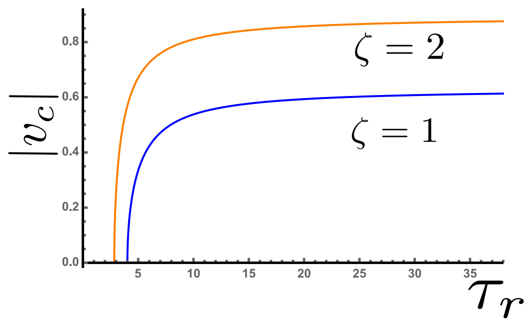


Figure T4: $v_c$ as a function of $\tau_r$ with $\ell = 1$, $f = 1$ and $\sigma = 1$. This demonstrates that as $\tau_r$ increases, $v_c$ goes to a constant value.

Unfortunately, the solution of the self-consistent equation cannot be obtained in a closed form even for a Gaussian forcing function. Therefore, we present its graphical solution in Fig. T3 for a Gaussian profile with $\sigma = 1$ for multiple $\tau_r$ and $\ell$, holding $\zeta = 1$ and $f = 1$ fixed. We explicitly calculate the scaling of $v_c$ with $\tau_r$ in Fig. T4 and demonstrate that it goes to a $\zeta$-dependent constant as $\tau_r$ increases. This is therefore consistent with $v_c$ being controlled by $\zeta$ with $\tau_r$ entering *only* through the dependence of $\zeta_c$ on it. This conclusion is reasonable in a biological context, since one cannot increase the speed of a cluster arbitrarily simply by increasing the correlation time of the medium of the substrate. The conclusion is also consistent with the results obtained in the experiments. Eq. (8) implied that for a delta-function, $v_c$ scales as $\zeta$ for large $\zeta$. We recover this result for a Gaussian profile with $\sigma \to 0$. However, for larger $\sigma$, this continuously changes with $v_c \sim \sqrt{\zeta}$ as we show in Fig. T5. In fact, for Gaussian profiles with $\sigma \approx 1$, the activity acts like temperature in a usual second-order phase transition, like in the Ising model. We show that $\zeta - \zeta_c$ scales as $v_c^2$ in Fig. T6 which implies that $v_c \sim \sqrt{\zeta - \zeta_c}$ as noted in the main text.

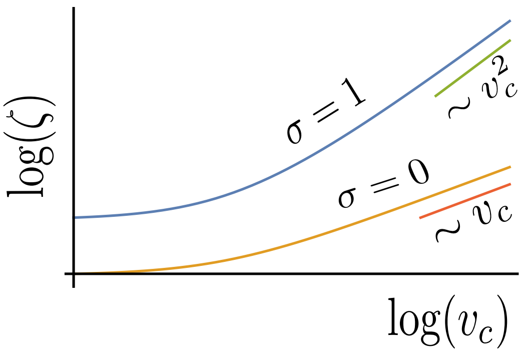


Figure T5: At large $v_c$, $\zeta \sim v_c^2$ for Gaussian distributions with $\sigma = 1$ and $\zeta \sim v_c$ for delta function distributions ($\sigma = 0$) for $f = 1$, $\ell = 1$ and $\tau_r = 1$. This implies that $v_c \sim \sqrt{\zeta}$ for large $\zeta$ for Gaussian profiles with $\sigma = 1$ in contrast to $v_c \sim \zeta$ for a $\delta$-function forcing.

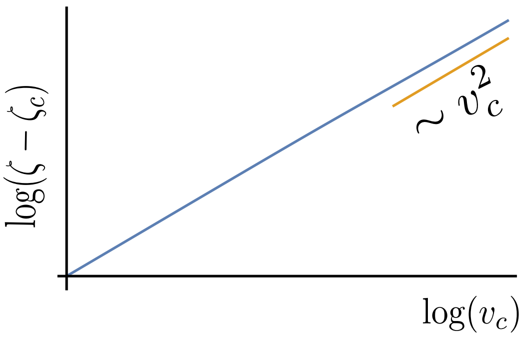


Figure T6: $\zeta - \zeta_c$ as a function of $v_c$ for $f = 1$, $\ell = 1$ and $\tau_r = 1$. This demonstrates that $v_c \sim \sqrt{\zeta - \zeta_c}$.

The fact that $v_c \sim \sqrt{\zeta - \zeta_c}$ and that $\tau_r$ and $\ell$ only affects $\zeta_c$ implies that we can heuristically view this as a phase transition controlled by a potential of the form $U = av^2/2 + bv^4/4$ with $a \propto \zeta_c - \zeta$. For $\zeta > \zeta_c$, i.e. for $a < 0$, the expectation value of $v$, $\langle v \rangle = v_c = \sqrt{-a/b} \propto \sqrt{\zeta - \zeta_c}$ and the potential has a double well with the minimum $U_{\text{min}} = -a^2/2b = -bv_c^2/2 \propto \zeta - \zeta_c$. This allows us to estimate the time before a motile cluster ($|v_c| > 0$) reverses its direction of motility (recall, that the direction of motility is determined by the sign of $v_c$) – within this heuristic potential picture, this corresponds to the crossing of a barrier of height $\Delta U = bv_c^2/2 \propto \zeta - \zeta_c$. Consider a noise of strength $2D$ that affects the dynamics of the cluster – Eq. (5) needs to be supplemented with a white noise with a variance $2D$ for this – the escape time from a potential of depth $\Delta U$ is then $\tau_e \propto e^{\Delta U/D}$. This implies that $\tau_e \sim e^{v_c^2} \sim e^{\zeta - \zeta_c}$. Therefore, at a constant value of $\zeta$, increasing $\tau_r$ decreases $\zeta_c$ and leads to an *increase* of $\tau_e$. However, $\tau_e$ goes to a constant for large $\tau_r$ (since $v_c$ goes to a constant as well). We have however assumed that the noise strength and the relaxation time $\tau_r$ of the medium can be tuned independently. This may not be the case for the experiments and a more accurate prediction of the escape time requires a more detailed description of the noise. However, it is gratifying that even this impressionistic description manges to reproduce the qualitative feature that increasing the relaxation time leads to an increase of the escape time and therefore, cell clusters move unidirectionally more persistently. Moreover, while the detailed scaling of $\tau_e$ with respect to $\zeta - \zeta_c$ depends on the form of the forcing function, the scaling with respect to $v_c$ holds irrespective of the forcing function. Since $v_c$ is expected to increase with $\tau_r$, at least for intermediate values of $\tau_r$, the persistence time of the cluster is expected to generally increase with $\tau_r$.

**Connection with a more detailed description of an isotropic motile layer**

To connect this simple picture of motility described in the previous section to a more detailed theory of a cluster described as an active droplet, we now describe the moving cluster by a phase field $\phi(\mathbf{x}, t)$, with $\phi = 1$ describing the interior of the cluster. The velocity field of the cluster is denoted by $\mathbf{v}(\mathbf{x}, t)$. The anisotropy of the the substrate is described by the apolar tensor $\mathsf{Q}(\mathbf{x}, t)$. The equation for the phase field is

$$\dot{\phi} + \nabla \cdot (\mathbf{v}\phi) = \nabla^2 \frac{\delta F}{\delta \phi} \tag{19}$$

where $F$ is the usual phase-field free energy. The overdamped equation for the velocity field is phenomenologically taken to be

$$\mathbf{v} = [\zeta_0 \mathsf{I} + \zeta_1 \mathsf{Q}] \cdot \nabla\phi - [M_0 \mathsf{I} + M_1 \mathsf{Q}] \cdot \phi \nabla \frac{\delta F}{\delta \phi} \tag{20}$$

where we have used an anisotropic mobility $[M_0 \mathsf{I} + M_1 \mathsf{Q}]$ with $\mathsf{I}$ being the identity tensor. $\zeta_0$ and $\zeta_1$ are active coefficients. The anisotropy of the substrate is affected by the motion as well as the shape of the cluster:

$$\dot{\mathsf{Q}} = -[\alpha - D\nabla^2]\mathsf{Q} + \lambda_0[\nabla\phi\nabla\phi - (1/2)(\nabla\phi)^2\mathsf{I}] + \lambda_1[\nabla\nabla\phi - (1/2)\nabla^2\phi\mathsf{I}] + \mu_0[\mathbf{v}\mathbf{v} - (1/2)|\mathbf{v}|^2\mathsf{I}] + \mu_1[\nabla\mathbf{v} + (\nabla\mathbf{v})^T - \nabla\cdot\mathbf{v}\mathsf{I}]. \tag{21}$$

The first direct term on the R.H.S. is the relaxation of the substrate when there is no aligning effect due to the cluster. The terms with the coefficients $\lambda_0$ and $\lambda_1$ describe the modification of the surface anisotropy due to the shape of the cluster (these terms can appear from free energy couplings $\mathsf{Q} : \nabla\phi\nabla\phi$ and $\mathsf{Q} : \nabla\nabla\phi$). The $\mu_1$ term describes usual flow-alignment. The term with the coefficient $\mu_0$ is more complicated. It exists only in systems on substrates but may be present *even* in passive systems where it can arise from a free energy term $\mathbf{Q} : \mathbf{v}\mathbf{v}$. We want to check whether the cluster can *spontaneously* break symmetry and acquire a non-zero average velocity $\langle \mathbf{v} \rangle$ in some direction we denote by $\hat{x}$. For simplicity, we also assume that all variations are only in this direction and reduce the two-dimensional problem to a one-dimensional one, i.e. $\phi \equiv \phi(x,t)$, $\mathbf{v} \equiv v_x(x,t)$. In this spirit, we assume that the apolar order parameter is

$$\mathbf{Q} = S(x) \begin{pmatrix} 1 & 0 \\ 0 & -1 \end{pmatrix} \tag{22}$$

This leads to the equations

$$\partial_t \phi + v_x \partial_x \phi = \partial_x^2 \frac{\delta F}{\delta \phi} \tag{23}$$

$$v_x = [\zeta_0 + \zeta_1 S]\partial_x \phi - [M_0 + M_1 S]\phi \partial_x \frac{\delta F}{\delta \phi} \tag{24}$$

$$\partial_t S = -[\alpha - D\partial_x^2]S + \frac{\lambda_0}{2}(\partial_x \phi)^2 + \frac{\lambda_1}{2}\partial_x^2 \phi + \frac{\mu_0}{2} v_x^2 + \mu_1 \partial_x v_x. \tag{25}$$

We now look at a drop which has fixed shape due to a high stiffness such that $\phi \approx 0$ outside a small region. We assume $\delta F/\delta \phi \approx 0$. We take $\phi$ to be $\approx 1$ between $L$ and $-L$ and 0 outside it. In this case, the average velocity of the cluster is

$$v_c = \frac{1}{2L}\int_{-L}^{L} v_x(x)dx. \tag{26}$$

This implies

$$v_c = \frac{\zeta_0}{2L}\int_{-L}^{L} \partial_x \phi dx + \frac{\zeta_1}{2L}\int_{-L}^{L} S(x)\partial_x \phi dx = \zeta_1 [S(x)\phi(x)]_{-L}^{L} - \frac{\zeta_1}{2L}\int_{-L}^{L} \phi(x)\partial_x S dx. \tag{27}$$

This implies

$$v_c = -\zeta_1 \frac{S(L) - S(-L)}{2L}. \tag{28}$$

In the limit $L \to 0$, $-\lim_{L\to 0} \zeta_1 S(L) - S(-L)/2L = -\zeta_1 \partial_x S|_{x_c}$. In the $S(x)$ equation, the velocity couplings lead to nonlinear terms in $S(x)$ or modify the coefficients $\lambda_0$ and $\lambda_1$. Therefore, the ordering is induced primarily by the terms $(\partial_x \phi)^2$ and $\partial_x^2 \phi$. Both of these terms are even functions of $x$ about $0$ for a $\phi$ profile that is an even function of $x$. These dynamical equations are equivalent to the ones we we considered in the last section and results in the same spontaneous symmetry-breaking transition mechanism discussed there. This demonstrates that a less impressionistic model of a cell-cluster reduces to the more impressionistic version of a point particle in a self-generated deformation field and justifies its consideration to provide a qualitative understanding of the experimental phenomenon.

#### Interpretation of the structural perturbation $S$ in the model

In the main text, we declared that the $S$ field can describe both a perturbation of the apolar organisation of the collagen filaments as well as the local density of the collagen. The first is obvious – $S$ in this case is the norm of the rank-2 apolar order parameter tensor and measures the local degree of apolar orientation. In the isotropic phase, the viscous relaxation dynamics of $S$ is generically given by Eq. 2[43]. However, the second requires more explanation not the least because density is a conserved variable and must be associated with a continuity equation. We now provide an interpretation of the equation of motion for $S$ to clarify this. Consider a slab geometry in which the collagen network of thickness $h$ rests on a solid substrate to which it is pinned. The cluster is on the top surface of the collagen network. We allow for only transverse displacements of the collagen network. Then, the overdamped dynamics of the displacement field $\mathbf{u}_\perp$ must be

$\gamma\partial_t \mathbf{u}_\perp = \lambda\nabla\mathbf{u}_\perp + \mu\nabla_\perp\nabla_\perp\cdot\mathbf{u}_\perp$ where $\gamma$ is a friction coefficient and $\lambda$ and $\mu$ are the shear and bulk modulus respectively ($\propto E$ in the main text). Note that $\mathbf{u}_\perp$ is a function of both the transverse, in plane variables, $x, y$ and the vertical one $z$ where $z$ is the coordinate that is measured from the bottom of the collagen layer with the network being pinned to the solid below it at $z = 0$ i.e., $\mathbf{u}_\perp(z = 0) = 0$. Therefore,

$$\gamma\partial_t \mathbf{u}_\perp = \lambda\nabla_\perp^2\mathbf{u}_\perp + \lambda\partial_z^2\mathbf{u}_\perp + \mu\nabla_\perp\nabla_\perp\cdot\mathbf{u}_\perp + \tilde{f}g(x - x_c)\delta(z - h) \tag{29}$$

where the last term denotes the forcing due to the cluster. We can now average this over the thickness of the network. This will generically yield

$$\gamma\partial_t \bar{\mathbf{u}}_\perp = \lambda\nabla_\perp^2\bar{\mathbf{u}}_\perp - \xi\frac{\lambda}{h^2}\bar{\mathbf{u}}_\perp + \mu\nabla_\perp\nabla_\perp\cdot\bar{\mathbf{u}}_\perp + \xi_2\tilde{f}hg(x - x_c) \tag{30}$$

where $\bar{\mathbf{u}}_\perp$ is the averaged displacement field and $\xi$ and $\xi_2$ are dimensionless numbers $> 0$ whose exact value depends on the displacement profile in $z$. It might appear surprising at first glance that while (29) is manifestly invariant under $\mathbf{u}_\perp \to \mathbf{u}_\perp + \text{const.}$, (30) is not – i.e., it loses *translation* invariance. This is a consequence of the pinning of the collagen layer to the solid at the bottom – this pinning fixes a preferred reference frame for the displacement field. Now, in the spirit of the one-dimensional model we allow for displacement only in $\hat{x}$ direction, i.e. only $\bar{u}_x$ is non-zero, and allow *that* to vary only along $x$, which yields

$$\gamma\partial_t \bar{u}_x = (\lambda + \mu)\partial_x^2\bar{u}_x - \xi\frac{\lambda}{h^2}\bar{u}_x + \xi_2\tilde{f}hg(x - x_c) \tag{31}$$

Identifying $\bar{u}_x$ with $S$, $\gamma h^2/(\xi\lambda)$ with $\tau_r$, $h^2(\lambda + \mu)/(\xi\lambda)$ with $\ell^2$ and $\xi_2\tilde{f}/\gamma$ with $f$, we can map this model onto (9). Since in a crosslinked gel, the dynamics of density fluctuations $\delta\rho$ is slaved to the displacement dynamics via $\partial_t(\delta\rho + \nabla_\perp\cdot\mathbf{u}_\perp) = 0$, which with the assumptions we have made can be rewritten as $\partial_t(\delta\rho + \partial_x \bar{u}_x) = 0$, this interpretation provides direct information about the density fluctuations as well.

Now we come to how the phenomenological coefficients in these equations are modified when the collagen gel gets more crosslinked. In the interpretation in which $S$ is the scalar orientational order parameter, crosslinking is likely to *reduce* $\tau_r$ while not significantly affecting $\ell$ since random crosslinking is likely to *hinder* orientational ordering. This, as discussed in the main text, leads to a *decrease* of persistent motion. In the interpretation in which $S$ is identified with the displacement field $\bar{u}_x$, $\tau_r = \gamma h^2/(\xi\lambda)$. Since crosslinking *increases* the stiffness of the network, which should *increase* the shear modulus, $\tau_r$ is again reduced and $\ell^2$, which is controlled by the ratio of bulk and shear moduli is not likely to be significantly affected. This implies that again in this interpretation, crosslinking should *reduce* the directed motion of the cluster.

### Supplementary Figures

Figure S1

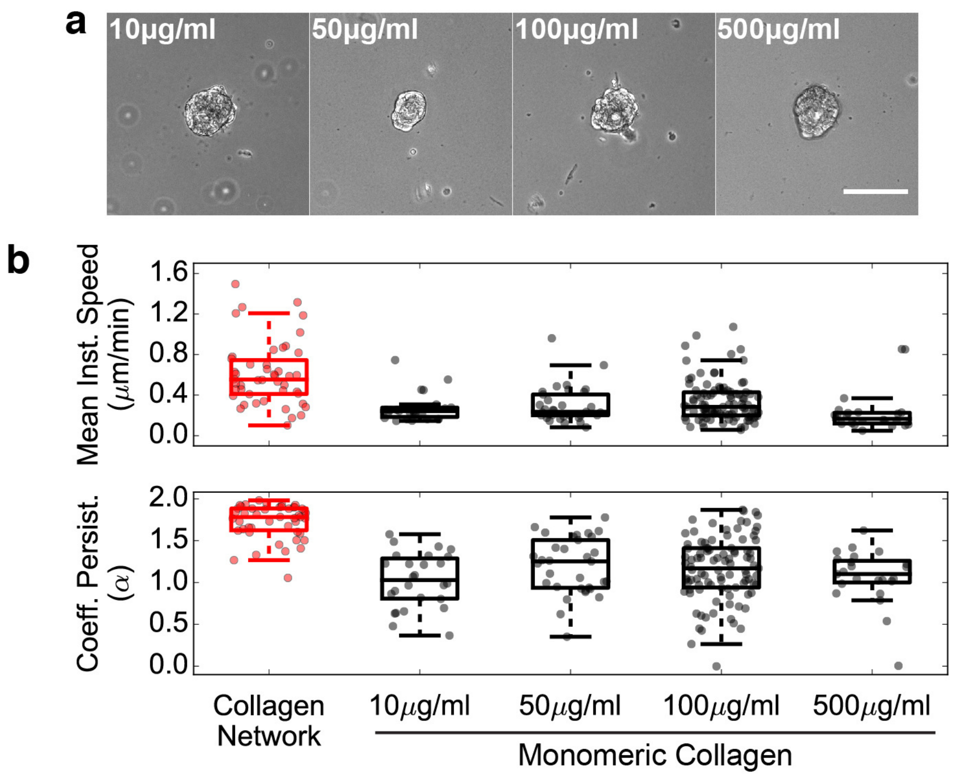


**Figure S1. Cluster migration on 2D gels does not depend on collagen concentration. a.** Example micrographs of cell clusters plated on PAA gels with different concentrations of monomeric collagen. Scale bar: 100$\mu$m. **b.** Boxplots of mean instantaneous speed and coefficient of persistence for PAA gels coated with a collagen network and different concentrations of monomeric collagen. Data represents n = 46, 26, 33, 97, 23 clusters from N = 5, 1, 1, 3, 1 independent experiments. Data for collagen network and 100$\mu$g/ml monomeric collagen is the same data as presented in Fig. 1c.

Figure S2

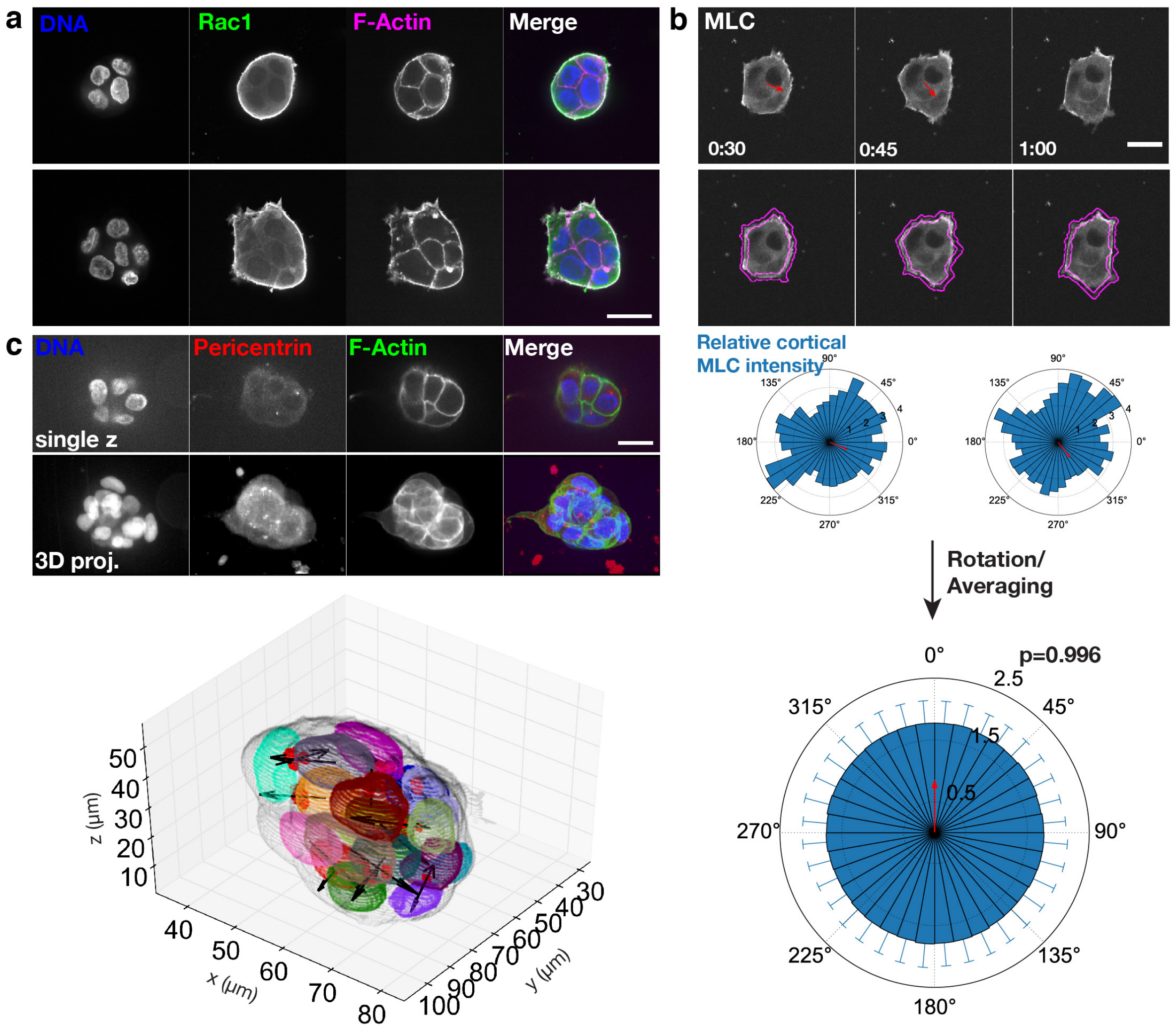


**Figure S2. Clusters exhibit supracellular cortex organization and do not display strong front-back polarity. a.** Fixed cell clusters plated on a collagen network and immunostained for Rac1 GTPase with DAPI (DNA) and Phalloidin (F-Actin). Shown are two representative examples from n = 21 clusters and N = 3 independent experiments. Scale bar: 25$\mu$m. **b.** Montage from timelapse of a myosin light chain (MLC)-GFP expressing A431 cell cluster. MLC-GFP intensity was measured in the peripheral cortical region (segmentation in magenta) around the perimeter of the cluster (see also Supplementary Movie 3). At each time point, the relative cortical intensity was measured for different angles around the perimeter (blue bars) as well as the cluster trajectory (red arrow). *Lower panel*: The relative cortical intensity of myosin (mean$\pm$SD), averaged over all time points for n = 39 clusters from N = 3 independent experiments. Scale bar: 25$\mu$m. p-value reflects Rayleigh test of uniformity. **c.** Example micrograph from cell cluster on a collagen network with centrosomes stained with pericentrin. Images displayed are a single $z$-slice and a 3D projection from the imaging stack. Scale bar: 20$\mu$m. *Lower panel*: 3D plot of the stained cluster in the upper panel, with the segmented cluster boundaries labeled in gray, segmented nuclei in multicolors and the paired centrosomes in red. Black arrows: centrosome orientation with respect to the nucleus center of mass. Representative example from n = 7 clusters from N = 1 independent experiment.

Figure S3

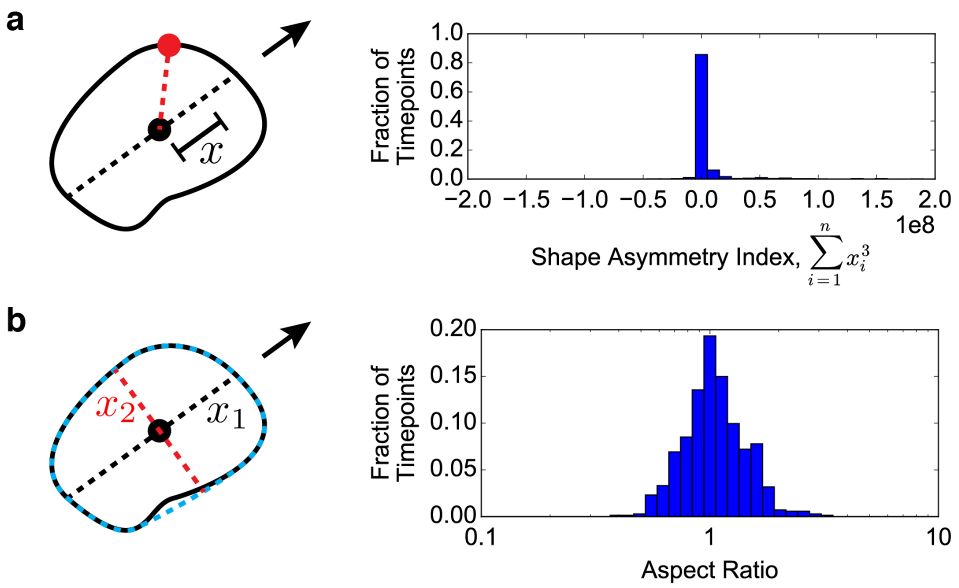


**Figure S3. Cluster shape is symmetric during migration. a.** Schematic of shape asymmetry quantification. A line (dotted black line) is drawn along the length of the cluster that is parallel with the migration direction (black arrow) and passes through the contour center of mass (black dot). At every pixel along the contour (for example, at the red dot), the distance, $x$, from the center of mass projected onto the center line is determined. The shape asymmetry index is calculated for each time frame as the sum of $x^3$ over all contour points. (Summing $x$ would always result in 0 by the definition of the center of mass, and summing $x^2$ would always be positive; therefore $x^3$ is summed.) *Right*: histogram of the sum of $x^3$ for all time points from n = 34 cells from N = 5 independent experiments. **b.** Schematic of aspect ratio quantification. The cluster length in the direction of migration, $x_1$, is determined from a line (dotted black line) that is parallel with the migration direction (black arrow) and passes through the contour center of mass (black dot). The cluster length perpendicular to the direction of migration, $x_2$, is determined from a line (dotted red line) that is perpendicular to the migration direction and passes through the contour center of mass. The aspect ratio is $x_1/x_2$. For determining $x_1$ and $x_2$, the Hull convex (blue dotted line) of the contour is used to avoid shape irregularities. *Right*: histogram of the cluster aspect ratio for all time points from n = 34 cells from N = 5 independent experiments.

Figure S4

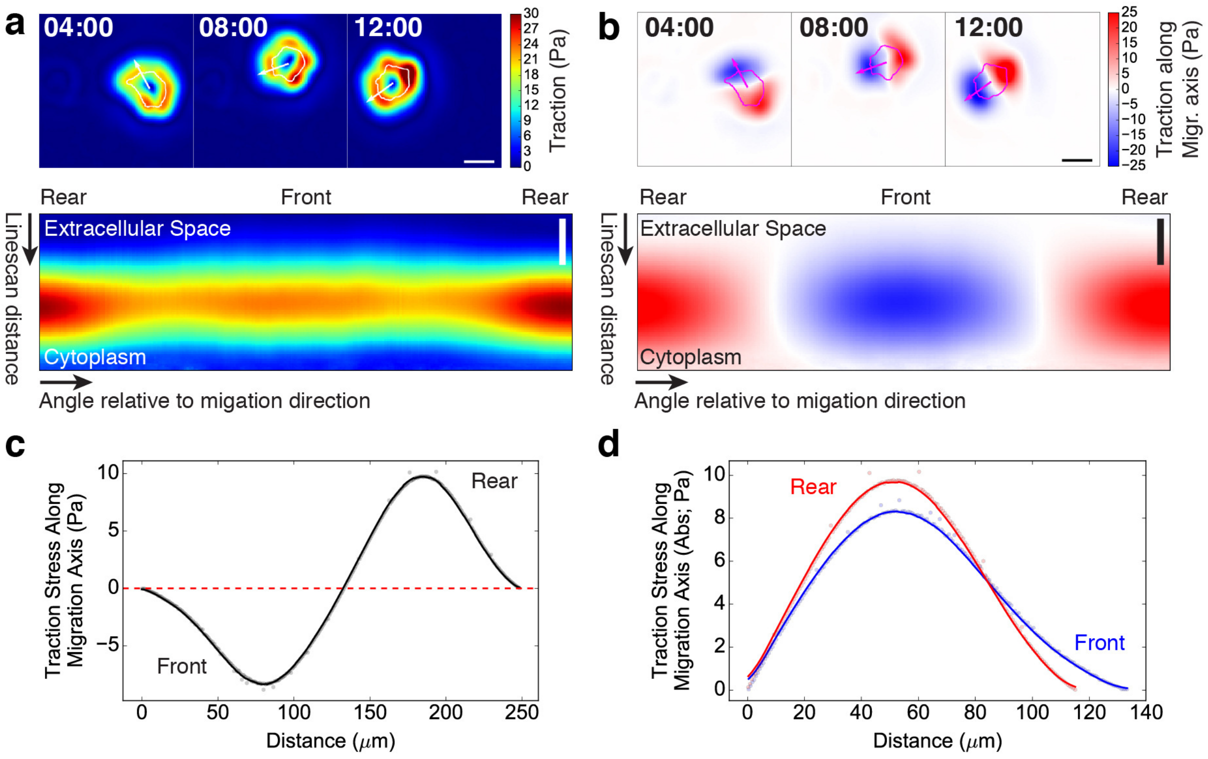


975 **Figure S4. Linescans of traction forces along the migration axis. a.** Magnitudes of traction forces exerted during migration by the cell cluster shown in Figure 1d. White line: cluster contour. White arrow: cell trajectory to the next time point. *Lower panel*: Unfolded radial linescans showing higher peak traction magnitudes in the front and rear of the cluster (mean over all timepoints). Horizontal axis: angle of the linescan with respect to the cluster front, ranging from -180° to 180° (the cluster front is at 0°). Vertical axis: distance along the linescan. Scale bar: 20$\mu$m. **b.** Traction forces projected along the axis of migration. Positive values indicate tractions in the same direction as the migration direction. Negative values indicate tractions in direction

opposite to the migration direction. Magenta line: cluster contour. Magenta arrow: cell trajectory to the next time point. The sum of the projected tractions is 0 at every time point due to force balance. *Lower panel*: Unfolded radial linescans of the tractions along the migration axis (mean over all timepoints). Axes as in *a, lower panel*. Scale bar: 20$\mu$m. **c.** Plot of a linescan of tractions along the migration axis from the front of the cluster to the rear of the cluster (averaged over all timepoints for the cluster shown in *a*,*b*). Blue dots: raw values. Blue line: smoothed data. The sum of the linescan is $\sim$0 due to force balance. **d.** Plot of the absolute value of tractions along the migration axis (from data shown in *c*). The linescan was cut in the middle (where the values crossed zero), and the front of the linescan was reversed in order to plot the front and back in the same direction. Dots: raw values. Lines: smoothed data.

Figure S5

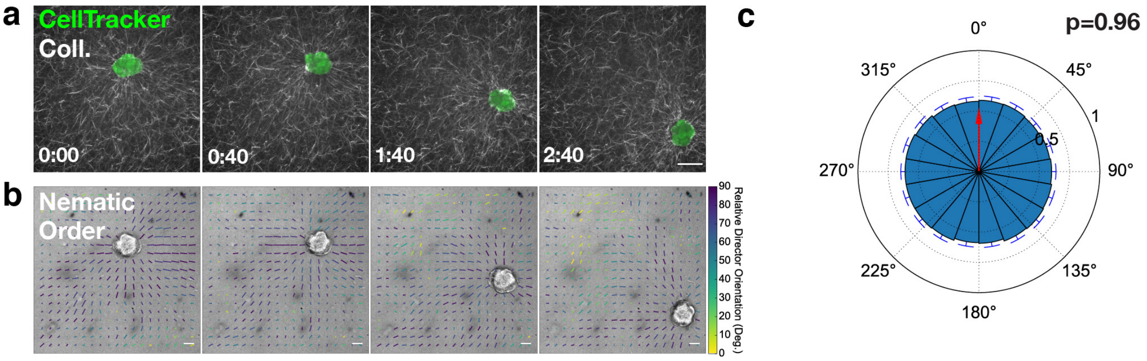


**Figure S5. Clusters align collagen networks into radially symmetric fibers. a.** Montage of a cluster migrating on a fluorescent collagen network (single $z$-slice on network surface; same images as in Fig. 2a). Scale Bar: 50$\mu$m. HH:MM. **b.** Brightfield montage of the cluster migrating in *a* overlaid with the local nematic order. Local nematic order is represented by rods whose direction indicates mean collagen fiber orientation (used as the nematic director, $\mathbf{n}$); rod color indicates the orientation of $\mathbf{n}$ with respect to the center of mass of the cluster (90°is oriented directly toward the center of mass, 0°is oriented away from the center of mass); rod length indicates the nematic order parameter, $S$ (scale rod: $S = 1$). **c.** Polar plot of the nematic order within 50$\mu$m of the cluster boundary (mean±SD) averaged over all time points for n = 28 cells from N = 3 independent experiments. p-value reflects Rayleigh test of uniformity.

### Supplementary Movies

**Supplementary Movie 1.** A431 cluster migrating on a 0.5kPa PAA gel coated with a thin collagen-1 network. Scale Bar: 100$\mu$m. HH:MM.

**Supplementary Movie 2.** A431 cluster migrating on a 0.5kPa PAA gel coated with 100$\mu$g/ml monomeric collagen-1. Scale Bar: 100$\mu$m. HH:MM.

**Supplementary Movie 3.** A431 myosin light chain (MLC)-GFP clusters migrating on collagen-1 networks. Magenta lines indicate region used for segmentation of the cluster cortex. Scale Bar: 50$\mu$m. HH:MM.

**Supplementary Movie 4.** A431 clusters migrating on a 0.5kPa PAA gel coated with a thin collagen-1 network analyzed by traction force microscopy. Arrows: *xy* traction stresses on the substrate. Scale Bar: 100$\mu$m. HH:MM

**Supplementary Movie 5.** A431 cluster migrating on a fluorescently-labeled collagen network. *Left*: the fluorescent collagen is shown in pseudocolor and the cell outline is shown in white. The white dot represents the cluster center of mass. Scale Bar: 50$\mu$m. HH:MM. *Right*: Separation of the segmentation into equal-length segments from front (yellow) to rear (purple).

**Supplementary Movie 6.** Rapid removal of an A431 cell cluster cultured on a fluorescent collagen network. *Left*: Brightfield. *Middle*: Collagen in grayscale with cluster segmentation in magenta. *Right*: Collagen in grayscale with vectors from PIV shown in colors according to angle. Time 0:00

refers to frame prior to addition of Trypsin/$NH_4OH$. Subsequent timestamps refer to time after Trypsin/$NH_4OH$ treatment. Scale Bar: 50$\mu$m. Scale Vector: 0.5$\mu$m/min. HH:MM.

**Supplementary Movie 7.** 3D displacement microscopy experiment for an A431 cell cluster cultured on a collagen network and rapidly removed using Trypsin/$NH_4OH$. Black arrows: *xy* displacement vectors. Color scale: *z* displacement vectors (negative values indicate displacement down toward the substrate). Time 0:00 refers to frame prior to addition of Trypsin/$NH_4OH$. Subsequent timestamps refer to time after Trypsin/$NH_4OH$ treatment. Scale Bar: 50$\mu$m. Scale Vector: 50$\mu$m. HH:MM.

**Supplementary Movie 8.** Rapid removal of an A431 cell cluster cultured on a fluorescent collagen network. *Left*: Brightfield. *Middle*: Collagen in grayscale with cluster segmentation in magenta. *Right*: Collagen in grayscale with vectors from PIV shown in colors according to angle. Time 0:00 refers to frame prior to addition of Trypsin/$NH_4OH$. Subsequent timestamps refer to time after Trypsin/$NH_4OH$ treatment. Scale Bar: 50$\mu$m. Scale Vector: 0.5$\mu$m/min. HH:MM.

**Supplementary Movie 9.** Rapid removal of an A431 cell cluster cultured on a fluorescent collagen network crosslinked with 1mM threose. *Left*: Brightfield. *Middle*: Collagen in grayscale with cluster segmentation in magenta. *Right*: Collagen in grayscale with vectors from PIV shown in colors according to angle. Time 0:00 refers to frame prior to addition of Trypsin/$NH_4OH$. Subsequent timestamps refer to time after Trypsin/$NH_4OH$ treatment. Scale Bar: 50$\mu$m. Scale Vector: 0.5$\mu$m/min. HH:MM.

**Supplementary Movie 10.** Rapid removal of an A431 cell cluster cultured on a fluorescent collagen network crosslinked with 10mM threose. *Left*: Brightfield. *Middle*: Collagen in grayscale with cluster segmentation in magenta. *Right*: Collagen in grayscale with vectors from PIV shown in colors according to angle. Time 0:00 refers to frame prior to addition of Trypsin/$NH_4OH$. Subsequent timestamps refer to time after Trypsin/$NH_4OH$ treatment. Scale Bar: 50$\mu$m. Scale Vector: 0.5$\mu$m/min. HH:MM.

**Supplementary Movie 11.** A431 cluster migrating on a fluorescently-labeled collagen network crosslinked with 1mM threose. *Left*: the fluorescent collagen is shown in pseudocolor and the cell outline is shown in white. The white dot represents the cluster center of mass. Scale Bar: 50$\mu$m. HH:MM. *Right*: Separation of the segmentation into equal-length segments from front (yellow) to rear (purple).

**Supplementary Movie 12.** A431 cluster migrating on a fluorescently-labeled collagen network crosslinked with 10mM threose. *Left*: the fluorescent collagen is shown in pseudocolor and the cell outline is shown in white. The white dot represents the cluster center of mass. Scale Bar: 50$\mu$m. HH:MM. *Right*: Separation of the segmentation into equal-length segments from front (yellow) to rear (purple).

**Supplementary Movie 13.** A431 cluster migrating on a collagen-1 network. Scale Bar: 100$\mu$m. HH:MM.

**Supplementary Movie 14.** A431 cluster migrating on a collagen-1 network crosslinked with 1mM threose. Scale Bar: 100$\mu$m. HH:MM.

**Supplementary Movie 15.** A431 cluster migrating on a collagen-1 network crosslinked with 10mM threose. Scale Bar: 100$\mu$m. HH:MM.

**Supplementary Movie 16.** Rapid removal of an individual A431 cell cultured on a fluorescent collagen network. *Left*: Brightfield. *Middle*: Collagen in grayscale with cluster segmentation in magenta. *Right*: Collagen in grayscale with vectors from PIV shown in colors according to angle. Time 0:00 refers to frame prior to addition of Trypsin/$NH_4OH$. Subsequent timestamps refer to time after Trypsin/$NH_4OH$ treatment. Scale Bar: 50$\mu$m. Scale Vector: 0.5$\mu$m/min. HH:MM.

**Supplementary Movie 17.** A431 single cell migrating on a fluorescently-labeled collagen network. *Left*: the fluorescent collagen is shown in pseudocolor and the cell outline is shown in white. The white dot represents the cluster center of mass. Scale Bar: 50$\mu$m. HH:MM. *Right*: Separation of the segmentation into equal-length segments from front (yellow) to rear (purple).

**Supplementary Movie 18.** A431 single cell migrating on a collagen-1 network. Scale Bar: 50$\mu$m. HH:MM.
